# Footprints of worldwide adaptation in structured populations of *D. melanogaster* through the expanded DEST 2.0 genomic resource

**DOI:** 10.1101/2024.11.10.622744

**Authors:** Joaquin C. B. Nunez, Marta Coronado-Zamora, Mathieu Gautier, Martin Kapun, Sonja Steindl, Lino Ometto, Katja M. Hoedjes, Julia Beets, R. Axel W. Wiberg, Giovanni R. Mazzeo, David J. Bass, Denys Radionov, Iryna Kozeretska, Mariia Zinchenko, Oleksandra Protsenko, Svitlana Serga, Cristina Amor-Jimenez, Sònia Casillas, Alejandro Sanchez-Gracia, Aleksandra Patenkovic, Amanda Glaser-Schmitt, Antonio Barbadilla, Antonio J. Buendia-Ruiz, Astra Clelia Bertelli, Balázs Kiss, Banu Sebnem Önder, Bélen Roldán Matrín, Bregje Wertheim, Candice Deschamps, Carlos E. Arboleda-Bustos, Carlos Tinedo, Christian Feller, Christian Schlötterer, Clancy Lawler, Claudia Fricke, Cristina P. Vieira, Cristina Vieira, Darren J. Obbard, Dorcas Orengo, Doris Vela, Eduardo Amat, Elgion Loreto, Envel Kerdaffrec, Esra Durmaz Mitchell, Eva Puerma, Fabian Staubach, Florencia Camus, Hervé Colinet, Jan Hrcek, Jesper G. Sørensen, Jessica Abbott, Joan Torro, John Parsch, Jorge Vieira, Jose Luis Olmo, Khalid Khfif, Krzysztof Wojciechowski, Lilian Madi-Ravazzi, Maaria Kankare, Mads F. Schou, Manolis Ladoukakis, Maria Josefa Gomez-Julian, Maria Luisa Espinosa-Jimenez, Maria Pilar Garcia Guerreiro, Maria-Eleni Parakatselaki, Marija Savic Veselinovic, Marija Tanaskovic, Marina Stamenkovic-Radak, Margot Paris, Marta Pascual, Michael G. Ritchie, Michel Rera, Mihailo Jelić, Mina Hojat Ansari, Mina Rakic, Miriam Merenciano, Natalia Hernandes, Nazar Gora, Nicolas Rode, Omar Rota-Stabelli, Paloma Sepulveda, Patricia Gibert, Pau Carazo, Pinar Kohlmeier, Priscilla A. Erickson, Renaud Vitalis, Roberto Torres, Sara Guirao-Rico, Sebastian E. Ramos-Onsins, Silvana Castillo, Tânia F. Paulo, Venera Tyukmaeva, Zahara Alonso, Vladimir Alatortsev, Elena Pasyukova, Dmitry Mukha, Dmitri Petrov, Paul Schmidt, Thomas Flatt, Alan O. Bergland, Josefa Gonzalez

## Abstract

Large scale genomic resources can place genetic variation into an ecologically informed context. To advance our understanding of the population genetics of the fruit fly *Drosophila melanogaster*, we present an expanded release of the community-generated population genomics resource *Drosophila Evolution over Space and Time* (DEST 2.0; https://dest.bio/). This release includes 530 high-quality pooled libraries from flies collected across six continents over more than a decade (2009-2021), most at multiple time points per year; 211 of these libraries are sequenced and shared here for the first time. We used this enhanced resource to elucidate several aspects of the species’ demographic history and identify novel signs of adaptation across spatial and temporal dimensions. We showed that patterns of secondary contact, originally characterized in North America, are replicated in South America and Australia. We also found that the spatial genetic structure of populations is stable over time, but that drift due to seasonal contractions of population size causes populations to diverge over time. We identified signals of adaptation that vary between continents in genomic regions associated with xenobiotic resistance, consistent with independent adaptation to common pesticides. Moreover, by analyzing samples collected during spring and fall across Europe, we provide new evidence for seasonal adaptation related to loci associated with pathogen response. Furthermore, we have also released an updated version of the DEST genome browser. This is a useful tool for studying spatio-temporal patterns of genetic variation in this classic model system.

## Introduction

*Drosophila melanogaster* is a foundational model system in biology. Seminal studies in this species have played important roles in the development of modern population genetics, from empirical tests of genetic drift to classic examples of adaptation (e.g., Buri 1956; Lewontin 1974; Parsons 1975; McDonald and Kreitman 1991; Powell 1997; Casillas and Barbadilla 2017; Flatt 2020). Beyond its role as a model genetic system (Hales et al. 2015), *D. melanogaster* has a fascinating natural history in its own right. The species originated in southern-central Africa (Lachaise et al. 1988; Lachaise and Silvain 2004; Sprengelmeyer et al. 2020), splitting from its sister taxon, *D. simulans,* between 1.4 and 3.6 million years ago (Obbard et al. 2009; Obbard et al. 2012; Suvorov et al. 2022). While the species may have originally been a marula fruit specialist in the seasonal woodlands of southern-central Africa (Mansourian et al. 2018; Sprengelmeyer et al. 2020), it later adapted as a human commensal, ultimately developing a cosmopolitan distribution across all human-inhabited continents (Kapun et al. 2021; Chen et al. 2024).

The recent development of genomic resources for *D. melanogaster* has led to key discoveries about its phylogeography. For example, demographic inference has revealed that modern fruit fly populations expanded out of Africa after the last glacial maximum ∼10,000 ya (Kapopoulou et al. 2020), entering Asia around 3-4 kya (Chen et al. 2024), and Europe around ∼1,800 ya (Sprengelmeyer et al. 2020). European populations split into spatially defined clusters across Europe ∼1,000 ya (Kapun et al. 2020; Kapun et al. 2021). In the past two centuries, African and European populations experienced a secondary contact event in North America and Australia, likely due to mercantile activities and immigration (Capy et al. 1986; David and Capy 1988; Caracristi 2003; Kao et al. 2015; Bergland et al. 2016). Unlike its sister species *D. simulans, D. melanogaster* is capable of overwintering across a broad swath of temperate habitats (Izquierdo 1991; Machado et al. 2016; but see Serga et al. 2015) and can establish resident populations across its range (e.g., Ives 1945; Ives 1970; Machado et al. 2016; Kapun et al. 2021; Nunez et al. 2024). In temperate regions, *D. melanogaster* reaches its largest local population size during the peak of the growing season (e.g., late summer and early fall) and drastically decreases upon the onset of winter. These yearly boom-and-bust cycles are responsible for estimates of “local” population size that are orders of magnitude smaller than the “global” population size (Duchen et al. 2013; Sprengelmeyer et al. 2020; Nunez et al. 2024).

Over the past two decades, *D. melanogaster* has been the subject of numerous population genomics studies, which have collectively illuminated our general understanding of the evolution, the demography and the genetic basis of adaptation (e.g., reviewed in Casillas and Barbadilla 2017; Haudry et al. 2020; Guirao-Rico and González 2021). Like many other cosmopolitan drosophilids, *D. melanogaster* populations commonly occur along spatially distributed environmental gradients (e.g., latitudinal and altitudinal) leading to the formation of clines, with a large body of work providing evidence for spatially varying (clinal) selection (De Jong and Bochdanovits 2003; Hoffmann and Weeks 2007; Fabian et al. 2012; Adrion et al. 2015; Mateo et al. 2018; Flatt 2020). Moreover, populations of *D. melanogaster* are known to experience strong fluctuating selection regimes across the changing seasons (e.g., Schmidt and Conde 2006; Bergland et al. 2014; Behrman et al. 2015; Rajpurohit et al. 2018; Erickson et al. 2020; Machado et al. 2021; Rudman et al. 2022; Nunez et al. 2024; reviewed in Johnson et al. 2023). For example, worldwide analyses of genetic variation have found that chromosomal inversion polymorphisms are often involved in clinal and/or seasonal adaptation (Lemeunier and Aulard 1992; Kapun et al. 2016; Kapun and Flatt 2019; Kapun et al. 2023; Nunez et al. 2024). Likewise, several studies have successfully linked clinally and/or seasonally varying polymorphisms in *D. melanogaster* to fitness-relevant phenotypes (Lemeunier and Aulard 1992; Schmidt et al. 2008; Cogni et al. 2014; Paaby et al. 2014; Kapun et al. 2016; Kapun et al. 2016; Durmaz et al. 2019; Kapun and Flatt 2019; Betancourt et al. 2021; Yu and Bergland 2022; Glaser-Schmitt et al. 2023; Kapun et al. 2023; Nunez et al. 2024). Populations of *D. melanogaster* can thus be thought of as powerful “natural laboratories” to study adaptation across spatial and temporal scales, and to disentangle the contributions of selection and demography (Jensen et al. 2005; Ometto et al. 2005; Teshima et al. 2006; Thornton and Jensen 2007; Pavlidis et al. 2010).

Despite the status of *D. melanogaster* as a model organism, generating genomic datasets that capture the breadth and depth of genetic and phenotypic variation across the cosmopolitan range of the species is a complex task for single research groups. Furthermore, existing data for this species are heterogeneous across studies: several studies use resequenced inbred lines (Langley et al. 2012; Mackay et al. 2012; Lack et al. 2015; Lack et al. 2016), while others use sequencing of outbred individuals sequenced as a pool (i.e., Pool-Seq; Schlötterer et al. 2014), and the two data types can be difficult to reconcile. For these reasons, we have previously developed the *Drosophila Evolution over Space and Time* (**DEST**; https://dest.bio/) resource, with the aim of facilitating collaborative population genomic studies in *D. melanogaster* (Kapun et al. 2021). The DEST resource is the result of the collaborative efforts of the European *Drosophila* Population Genomics Consortium (**DrosEU**, https://droseu.net/; Kapun et al. 2020) and the *Drosophila* Real-Time Evolution Consortium, DrosRTEC (Machado et al. 2021). DEST represents both a tool for mapping genomic data, as well as an open-access data repository of worldwide genetic variation in the fruit fly. As a bioinformatics tool, DEST is a pipeline for mapping Pool-Seq reads to a hologenome reference of fly (i.e., *D. simulans* and *D. melanogaster*) and microbial genomes, as well as for removing contamination from other species, such as *D. simulans.* The tool is a highly modular mapping pipeline that uses a Docker image (Boettiger 2015) and *Snakemake* (Köster and Rahmann 2012) to ensure independence of operating systems. As a genomic panel, the original release of the dataset (DEST 1.0) consisted of 271 Pool-Seq *D. melanogaster* samples (> 13,000 flies) collected in more than 20 countries on four continents at different seasons and across multiple years. Using these data, we had previously described general patterns of phylogeographic structure across four continents, developed a panel of geographically informative markers (**GIMs**) to assess the provenance of fly samples with 90% accuracy, and we applied demographic inference tools (Jouganous et al. 2017) to infer the history of population subdivision in Europe (Kapun et al. 2020).

Here, we introduce the second release of the DEST resource (DEST 2.0), with substantial expansions in several methodological and biological aspects. From a methodological perspective, we have broadened the utility of our Docker application to allow for single end-reads to be mapped, a change that allows for older datasets to be integrated into DEST. We have explored levels of contamination by other species in DEST pools using a new highly efficient *k*-mer based approach (Gautier 2023). We have also estimated genome-wide rates of recombination using our Pool-Seq data by applying a deep learning approach (*ReLERNN*; Adrion et al. 2020). All data on genetic variation and population genetic summary statistics can be visualized and retrieved using our new and improved genome browser, which has been built with the latest JBrowse version 2 (Diesh et al. 2023).

From a biological standpoint, DEST 2.0 includes a substantial expansion of the size and scope of the initial dataset. The current release includes 530 high quality Pool-Seq samples (>32,000 flies), comprising a combination of the previous DEST release with newly sequenced pools, collected between 2016 and 2021 by DrosEU, as well as publicly available Pool-Seq samples from published studies of wild-derived *D. melanogaster* (Reinhardt et al. 2014; Svetec et al. 2016; Fournier-Level et al. 2019; Lange et al. 2022; Nunez et al. 2024). To showcase the utility of DEST 2.0, we performed several analyses to infer demography and selection, powered by the rich spatial and temporal density of our dataset. Below, we divide these analyses into two general categories: “*spatial insights*” and “*temporal insights*”. For each category, we highlight case studies of demographic inference and genome-wide scans for adaptive differentiation. Our analyses provide novel insights into patterns of demography and selection of natural *D. melanogaster* populations and generate hypotheses that can be tested with the power of the *Drosophila* genetics toolbox in future work. In general, our work illustrates the value of DEST 2.0 as an open resource for the *Drosophila* evolutionary genetics and genomics community.

## Results

### DEST 2.0, an expanded *Drosophila* population genomics resource

The current DEST release (v2.0) includes 530 high-quality samples as well as an additional 207 pools of varying quality (excluded from the analysis; see **Table S1**). In its totality, the 737 pooled libraries originated from multiple sources including both releases of the DEST dataset (i.e., v1.0 and v2.0), the *Drosophila* Genome Nexus (**DGN**; Lack et al. 2016; including one sample from *D. simulans*), as well as from previous publications (i.e., Reinhardt et al. 2014; Svetec et al. 2016; Fournier-Level et al. 2019; Lange et al. 2022; Nunez et al. 2024). The 737 samples within DEST 2.0 vary in sequencing characteristics, ranging from a read depth (abbreviated as “**RD**”) of 4X to 300X and from an effective haploid sample size (**n_e_**; the sample size accounting for pool size and pool-seq sampling effects) of 3.7 to 77.2 (**Fig. S1**; see **Text S1**; Kolaczkowski et al. 2011; Feder et al. 2012; Gautier et al. 2013). To ensure the highest possible quality of each sample, we calculated a battery of sequencing statistics including rate of PCR duplication, fraction of missing data, coverage, and number of private single nucleotide polymorphisms (SNPs) across the totality of the dataset (all 737 pools). In addition, we also estimated the pN/pS statistic (i.e., the ratio of the number of genome-wide non-synonymous polymorphisms to the number of genome-wide synonymous polymorphisms, as in Kapun et al. 2021; **Fig. S2**), and assessed non-*D. melanogaster* contamination through competitive mapping and k-mer approaches (Kapun et al. 2021, Gautier 2023; **Fig. S3**). Next, we used a principal component analysis (PCA) on all quality control metrics to assess whether samples should be included or excluded from downstream analyses (see **Fig. 2A** and **Fig. S4**; see Materials and Methods: *Estimation of nucleotide diversity*). Finally, 136 samples that consisted of multiple replicates from the same locality each with low coverage were collapsed into a single sample. For a more detailed description on Data filtering procedures and recommendations for users see **Text S2**. Based on the results of these analyses, we obtained a final high-quality dataset of 530 samples and 4,789,696 SNPs, across autosomes and the X chromosome for downstream analyses. The high quality dataset contains representative samples from 45 countries across all continents (22 from Africa, 40 from Asia, 302 from Europe, 141 from North America, 17 from Australia, and seven from South America; **Fig. 1A**) and across a time span of 12 years (2009-2021). In total, our 530 high-quality samples represent 164 localities, of these, 112 were sampled only in one year (68%), 18 were sampled across two years (11%), and the rest (34; 21%) were sampled multiple times across several years (**Fig. 1B**). Overall, descriptions and basic subsetting of SNP statistics for DEST 2.0 are shown in Table 1. Unless stated otherwise, all of the following analyses are based on the 530 high-quality samples.

**Figure 1.**
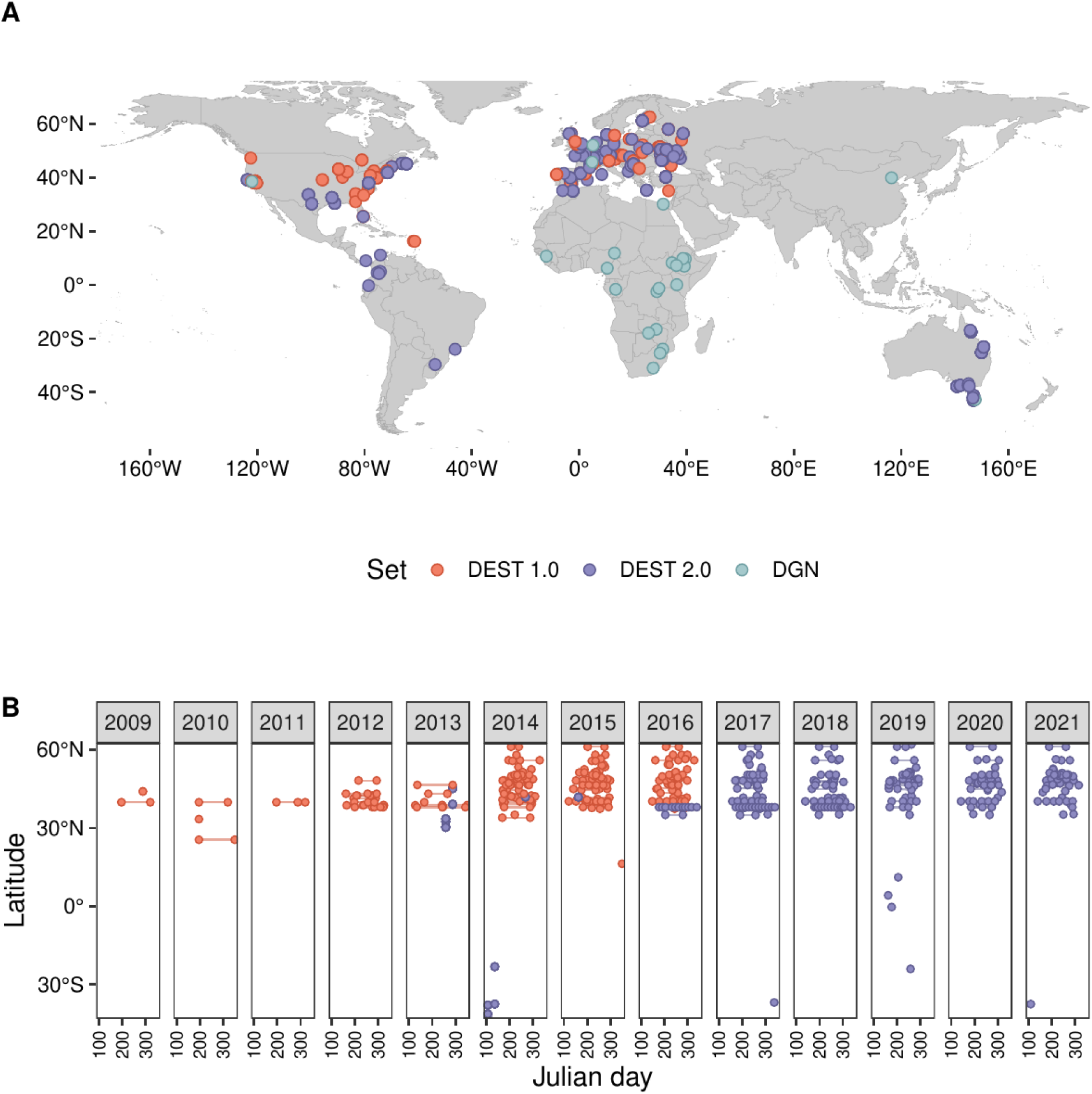
Spatial and temporal scales of DEST. **(A)** World map showing samples part of DEST 1.0 (Kapun et al. 2020), DEST 2.0 (this study), and the DGN (Lack et al. 2016). **(B)** Sampling density across a decade of sampling contained in the DEST dataset. The colors are consistent with panel A.

**Figure 2.**
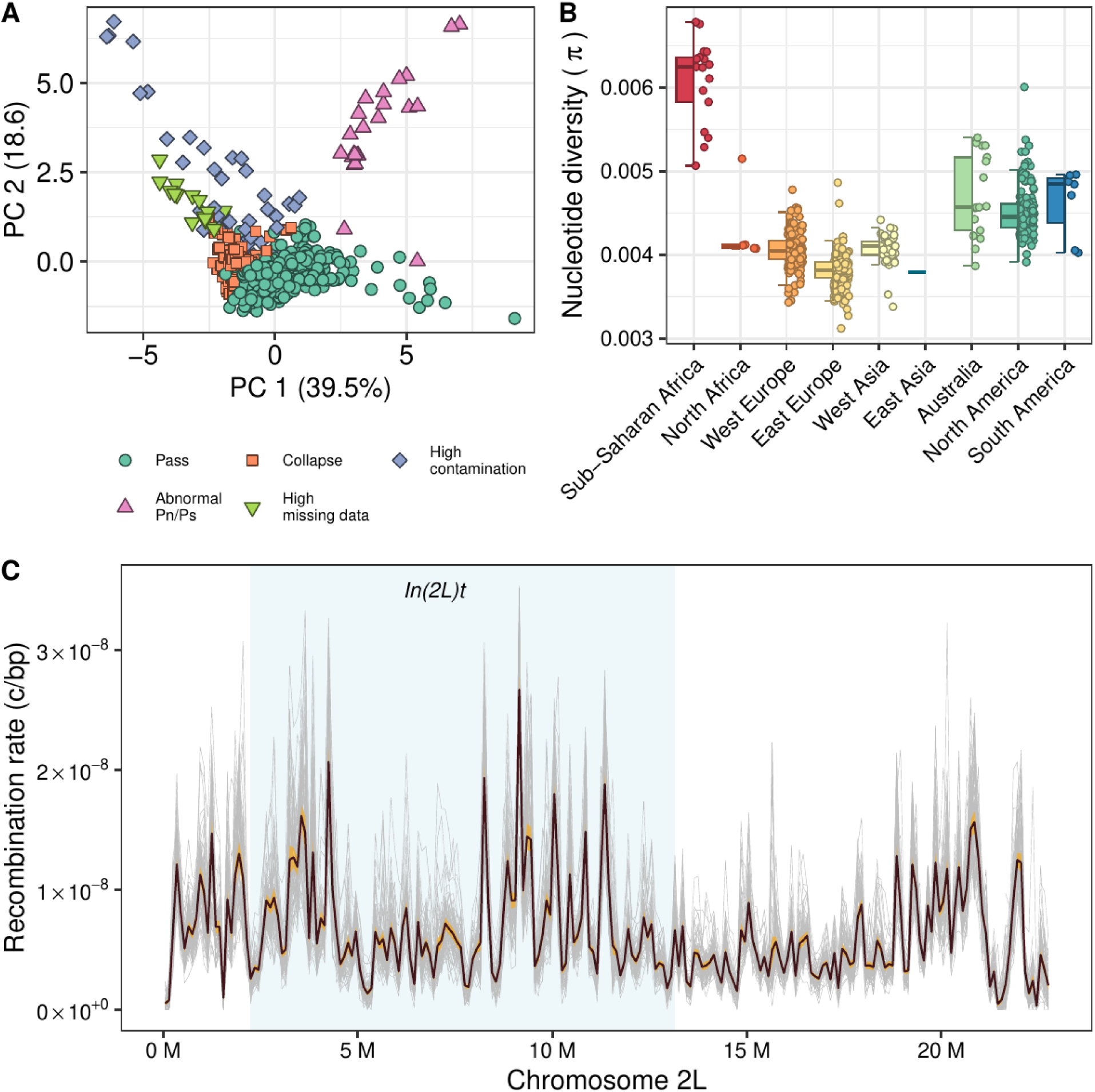
Patterns of filtering, genetic variation, and recombination in DEST 2.0. **(A)** Visualization of filtering information of samples using PCA. Each dot is a sample’s QC metric and the color indicates the filtering decision (legend: Pass: samples that pass filter and are used in downstream analyses; Collapse: biological and/or technical replicates collapsed into a single representative sample; otherwise samples were excluded due to abnormal pN/pS levels of high levels of missing data or contamination). **(B)** Nucleotide diversity (*π*) calculated across continents (see *Estimation of nucleotide diversity* for details). **(C)** Recombination landscape of chromosome 2L in samples representative of the 75 *D. melanogaster* populations analyzed (one gray line per sample). Light blue area highlights the region spanning the *ln(2L)t* inversion. Average (black line) and overall distribution envelope (orange shaded ribbon; delineated by the average values +/- 1.96 s.d.) are shown.

**Table 1:** SNP calling information for DEST 2.0 across major autosomes and chromosome X. SNPs inside the inversion are estimated of *In(2L)t* for *2L*, *In(2R)NS* for *2R*, *In(3L)P* for *3L*, and the joint region among *In(3R)K, In(3R)P,* and *In(3R)Mo*. Estimated recombination rates (i.e., rate of cross-over; “*c*”). Functional annotations are only reported for biallelic sites.

| SNP type | 2L | 2R | 3L | 3R | X |
| --- | --- | --- | --- | --- | --- |
| Total (All) | 1,080,586 | 901,878 | 1,069,441 | 1,212,752 | 525,039 |
| Bi-allelic | 1,048,510 | 877,852 | 1,039,460 | 1,182,310 | 516,077 |
| Inside inversions | 569,713 | 228,826 | 631,556 | 159,598 | NA |
| In recombining regions ($c > 0$) | 997,162 | 836,457 | 976,915 | 1,074,768 | 482,162 |
| Protein-coding | 796,420 | 731,794 | 793,866 | 944,372 | 40,4881 |
| Intergenic | 828,039 | 659,966 | 824,903 | 929,539 | 401,586 |
| Synonymous | 95,275 | 91,052 | 90,635 | 101,504 | 49,055 |
| Non-synonymous | 71,534 | 75,921 | 72,843 | 90,905 | 25,072 |
| Proportion of missing data | 0.0511 | 0.0507 | 0.0508 | 0.0493 | 0.0533 |

### Estimates of nucleotide diversity and recombination rates

To describe patterns of genetic variation in the DEST 2.0 data, we analyzed nucleotide diversity *π* (Tajima 1983; Tajima 1989) estimated with *npStat* (Ferretti et al. 2013). As previously observed (e.g., Begun and Aquadro 1993; Andolfatto 2001; Mackay et al. 2012; Kapun et al. 2021), we found that sub-Saharan African populations had higher levels of genetic variation than other populations (**Fig. 2B**), consistent with out-of-Africa demography (Li and Stephan 2006; Lack et al. 2016; Arguello et al. 2019; Kapopoulou et al. 2020; Kapun et al. 2021).

We inferred levels of genome-wide recombination across 75 samples representative of the populations analyzed (see Materials and Methods: *Recombination landscape*) using the deep learning method *ReLERNN* (Adrion et al. 2020; see **Fig. 2C; Fig. S5**). Overall, recombination rate is highly heterogeneous among samples and, among chromosomes (two-way ANOVA, *F*_74,296_ = 20.0, *P* < 1.0×10^-25^, and *F*_4,296_ = 1605.1, *P* < 1.0×10^-25^, respectively; Tukey’s HSD tests, all pairwise comparisons between chromosomes *P* < 1.0×10^-7^, except for 3R *vs*. 2R, where *P* = 0.073). In most populations there is a statistically significant positive correlation between recombination rate and genetic diversity, consistent with recurrent genetic hitchhiking and background selection (Begun and Aquadro 1993; **Table S2**).

The presence of common cosmopolitan inversions had a noticeable impact on the recombination landscape. Average recombination rates were significantly lower around the inversion breakpoints for five out of the seven inversions analyzed (Wilcoxon test, *P* < 0.01; for inversions *In(2L)t*, *In(3L)P*, *In(3R)Payne*, *In(3R)C* and *In(3R)K*; **Table S3**). Recombination was also lower for those regions spanning the three inversions than for the rest of the chromosome (Wilcoxon test, *P* < 0.01; for inversions *In(2R)NS*, *In(3R)Payne* and *In(3R)K*; **Table S3**).

PCA analyses showed that populations belonging to the same geographic cluster share similar recombination landscapes (**Fig. S6**; see **Table S1** for metadata). The geographic clustering is more evident when considering relative values of recombination, i.e., the ratio of the average recombination rate of each window to the average recombination across the respective chromosome, and is therefore informative on the recombination landscape rather than the absolute recombination rate (compare panels A and B with panels C and D in **Fig. S6**).

### Spatial population structure is defined by latitudinal and longitudinal clines

To investigate patterns of population structure in the DEST 2.0 dataset, we performed PCA on all 530 samples that passed quality filters. We used biallelic SNPs from the euchromatic regions of the four major autosome arms (**Figs. 3A-B**; also see **Fig. S7**). When all autosomes are considered, PC1 divides samples from sub-Saharan Africa from all other continents. At the level of individual regions, PC1 is correlated with both latitude and longitude in North America (*r* = -0.7; *P* = 2×10^-16^ and *r* = -0.59; *P* = 2.2×10^-16^, respectively) and longitude in Europe (*r* = -0.46; *P* = 2.2×10^-16^; **Fig. 3C-D**). These patterns of population structure were consistent with previously published studies (Kapun et al. 2020; Kapun et al. 2021; Machado et al. 2021). Both PC1 and PC2 primarily divided African samples from all other clusters, and PC2 also separated samples in Europe from samples in North America, South America, and Australia. PC3 primarily resolved discrete European clusters and also suggests that North American, South American and Australian samples behave like admixed samples (Ma and Amos 2012).

**Figure 3.**
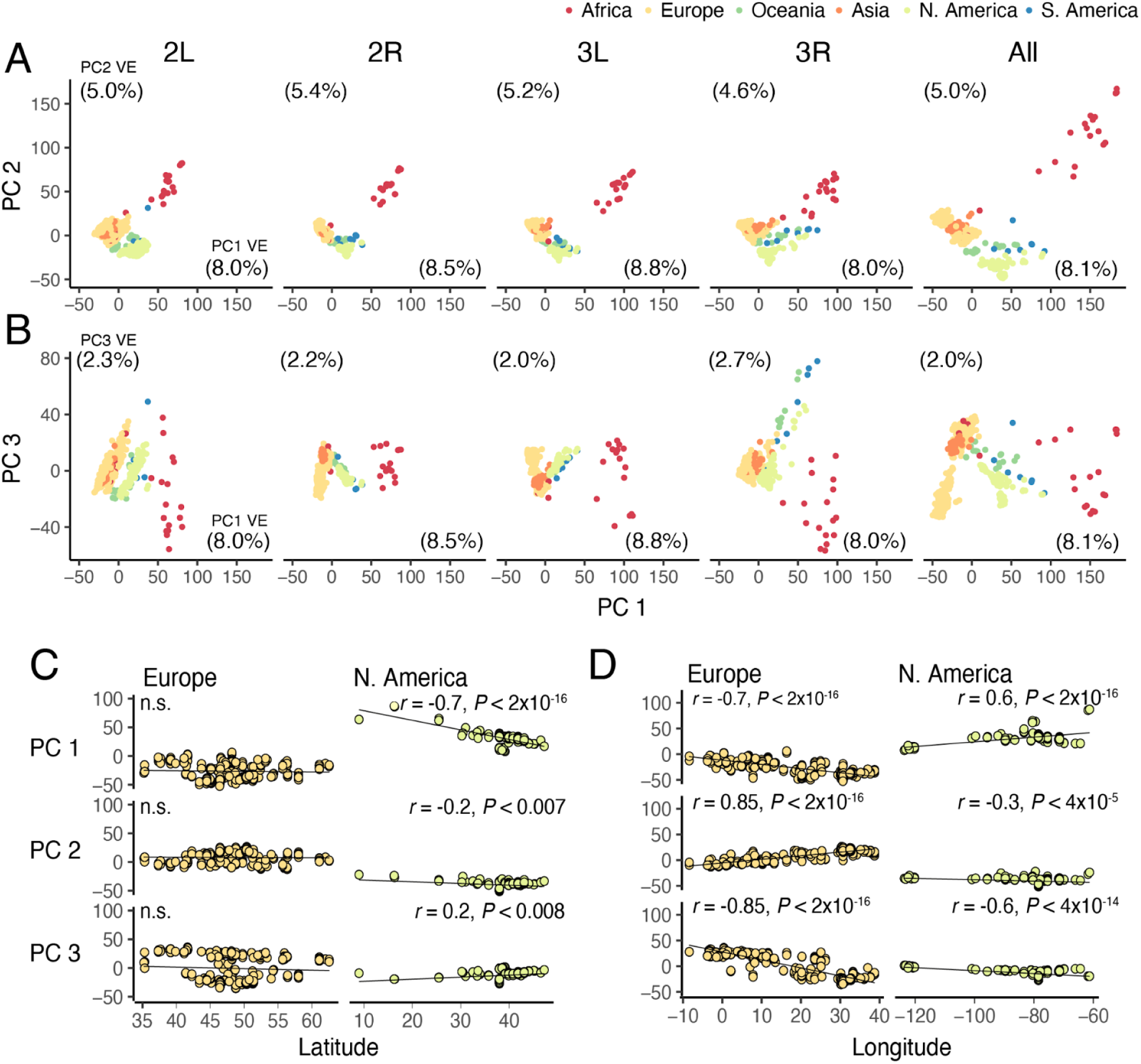
Principal component analysis and projections. **(A)** PCA projections showing PCs 1 and 2. Analyses were done for each chromosome arm and all arms combined. The proportion of variance explained (VE) is shown at the corners of each axis. (**B)** PCA projections showing PCs 1 and 3. (**C)** Projections of PCs 1, 2, and 3 relative to latitude for Europe and North American pools. (**D)** Same as C but for longitude. Notice that, in this analysis, Asia refers primarily to samples from Turkey (which is located in Western Asia).

The patterns seen across chromosome-specific PCA were strongly correlated to that of the whole genome for both PCs 1 and 2 (*r*_2L-All_ = ∼0.97, *r*_2R-All_ = ∼0.98, *r*_3L-All_ = ∼0.97, *r*_3L-All_ = ∼0.96; note that all *P* are < 1.0×10^-15^). PC3 is peculiar in that the whole-genome results were similar only to those for chromosomes 2R (*r*_2R-All_ = 0.95; *P* = 2.2×10^-16^) and 3L (*r*_3L-All_ = -0.95; *P*-value = 2.2×10^-16^), but not for 2L (*r*_2L-All_ = 0.18; *P* = 1.4×10^-5^) or 3R (*r*_3R-All_ = 0.05; *P* = 0.17). This observation suggests that the signal captured by PC3 at 2L and 3R were strongly influenced by the frequencies of *In(2L)t* and *In(3R)Payne*, two large adaptive cosmopolitan inversion polymorphisms (e.g., Kapun et al. 2023; Nunez et al. 2024).

We investigated clines in the frequencies of cosmopolitan inversion polymorphisms in DEST 2.0 using inversion-specific SNPs that are in strong linkage disequilibrium with the inversion breakpoints (Kapun et al. 2014; **Fig. S8**). Many inversions showed significant clinal patterns along latitude or longitude that were consistent across different continents (see **Table S4** for statistical details). Our results are in line with previous observations, in particular for *In(3R)Payne* (Lemeunier and Aulard 1992; Kapun et al. 2016; Kapun and Flatt 2019; Kapun et al. 2020; Kapun et al. 2023), which showed significant latitudinal clines in North America, Europe and along the Australian east coast. Notably, these patterns did not differ across sampling years in Europe and Australia, indicating temporal stability of the clines on these continents. Latitudinal clines were also significant for *In(2L)t* and *In(3R)Mo* in North America and Australia, and for *In(2R)NS* and *In(3L)P* in North America, Australia and Europe. Additionally, while overall not being very frequent, *In(2R)NS* exhibited a highly significant longitudinal cline across European populations.

### Characterizing latent population structure in European and North American populations

We applied *k*-means clustering analysis on the first three autosomal PCs to identify spatially defined clusters. First, with *k* = 4 clusters we fully recapitulated the results of DEST 1.0 (**Fig. 4A**), with clusters composed of sub-Saharan African samples, the Americas, and two clusters in Europe (as in Kapun et al. 2021; Europe West [**EU-W**] and Europe East [**EU-E**]). North African and West Asian samples clustered with EU-W. Australian samples were split between the clusters dominated by Western Europe and the Americas. We also estimated population clusters using *k* = 8, which was estimated to be the optimal value based on the gap statistic (Tibshirani et al. 2001; **Fig. 4B-inset**). For *k* = 8, new hypotheses of latent structure emerged (**Fig. 4B**). In Europe, the previously known EU-W and EU-E clusters appeared, separated by a putative third cluster at the boundary between EU-E and EU-W (i.e., an “overlapping zone”; **Fig. 4C**). Newer populations (namely the Americas and Australia), previously dominated by a single cluster, were divided into three clusters: the Caribbean and most of South America (henceforth “Latin America”), a southeast U.S. coastal group (henceforth “Southeast”), and all other samples from the Americas (henceforth “mainland”; see green, yellow, and pink points, respectively, in **Fig. 4B**). Notably, samples from Australia do not show any new levels of clustering when *k* = 8, relative to *k* = 4. Instead, they retain their original cluster association, whereby samples from the south of the continent cluster with samples from EU-W, and those from the north cluster with North American populations (**Fig. 4A** and **4B**). We used model-based demographic inference with *moments* (Jouganous et al. 2017) to test the statistical support of these additional populations suggested by the *k* = 8 analysis while simultaneously estimating demographic parameters. Specifically, we fit simple, neutral population history models that we call “one-population,” “split,” “admixture,” and “two-splits” (see **Fig. S9**; see description in the Materials and Methods: *Demographic inference with moments*) to subsets of the DEST 2.0 variant data consisting of the Southeast and mainland clusters, all samples from the Americas, and European samples (**Table S5**).

**Figure 4:**
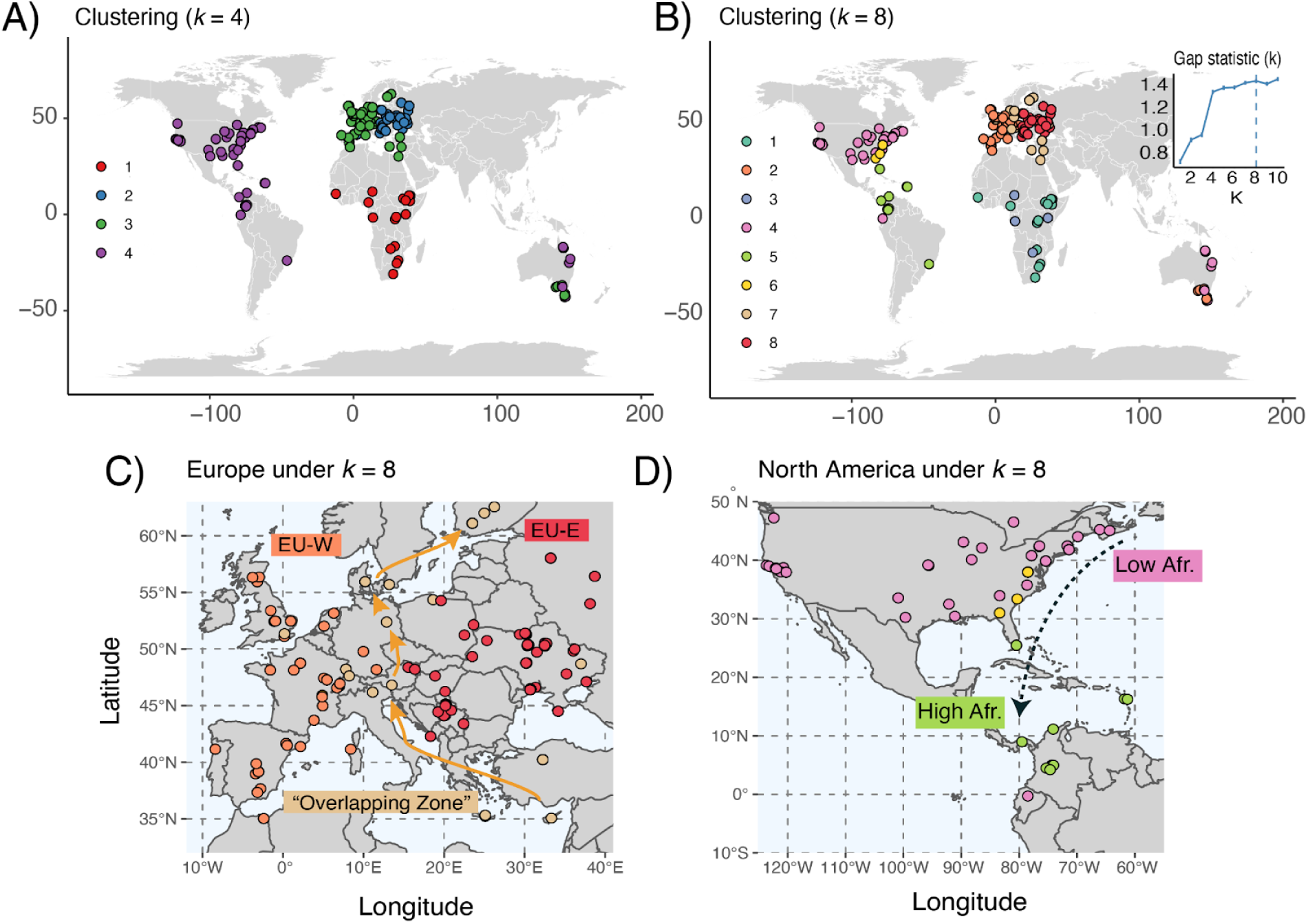
Spatial population structure and admixture in worldwide *Drosophila*. **(A)** Clustering map, based on PCA projections 1-3 built using *k* = 4 (as reported in DEST 1.0). **(B)** Same as A but with *k*=8 (the optimal number of clusters as defined by a heuristic Gap statistic search). **(C)** Zoom view of *k* = 8 into Europe to show the hypothetical overlap zone. **(D)** Zoom view of *k* = 8 into North America showing the hypothetical “Latin America” cluster (green) and Southeast cluster (yellow).

First, we fit the “one-population” and two-population “split” models to the Southeast and mainland clusters in North America to conclude that “one-population” better describes the region (Wilcoxon signed-rank test on model likelihoods, *P* = 7.02 x 10^-7^; **Fig. S10A**). This result, in which there is no strong evidence of historic divergence between the two clusters, along with low *F*_ST_ (0.034), supports the parsimony of clustering at *k* = 4. Thus, it is likely that the primary cause of the Southeast cluster in *k* = 8 analysis is the disproportionately dense sampling around Charlottesville, VA.

We then fit the “one-population” and “split” models to a population consisting of the Southeast and mainland clusters and the Latin America cluster, concluding again that “one-population” outperforms “split” (Wilcoxon signed-rank test on model likelihoods, *P* = 6.90 x 10^-9^; **Fig. S10B**). This result is complemented by the low *F*_ST_ = 0.062. This secondary result supports prior treatment of all flies of the Americas as a single cluster. This result does not contradict our findings of clines within the Americas, because the *demes*-type models employed rely on discretizing geography, and are thus largely blind to gradual changes with location.

In Europe, we conducted model comparisons among a two-population “split” model, three variants of the three-population “admixture” model (in which EU-W, the overlap region, and EU-E are respectively treated as the admixed population), and three variants of the three-population “two-splits” model (in which EU-W, the overlap region, and EU-E are respectively treated as a sister group to the other two populations). As in the Americas, we found support for the parsimonious two-population models that does not include the overlap zone as a discrete population (corrected Dunn’s tests on model likelihoods, *P* = 3.3 x 10^-7^; **Fig. S10C**). This result and the low three-way *F*_ST_ (0.036), indicate that only the EU-E and EU-W clusters are distinguished as discrete populations, and that the overlap zone may simply be an active area of gene flow between EU-W and EU-E. Overall, these findings suggest that the optimal demographic partitioning of the data coincides with clustering at *k* = 4, as reported in the original DEST release.

Next, we investigated the signals in the data that may have given rise to the clusters proposed by *k* = 8. We focused our analyses on the role of African–European admixture in the samples, as this is a primary driver of standing genetic variation in recently expanded populations (Bergland et al. 2016). To accomplish this, we first modeled the proportion of African and European admixture in the Americas and Australian pools as a linear combination of two “ancestral populations” from Europe and Africa (see **Dataset S1**). Our estimates of African admixture were consistent with previously published results (i.e., a positive, albeit non-significant, correlation between African admixture and latitude in Australia, β_African anc._ = 0.003, *P* = 0.162, see **Fig. 5A**; and a significant negative pattern in North America, β_African anc._ = -0.005, *P* = 2.5×10^-22^, see **Fig. 5B**; Bergland et al. 2016). We calculated these estimates in the newly collected samples from South America and observed a trend of increasing African ancestry near the equator (β_African anc._ is 0.002, *P* = 0.139, **Fig. 5C**). We also estimated the relationship between levels of admixture and longitude in North America. Here, we identified a significant association between longitude and ancestry (LM; β_African anc._ = 0.0014, *P* = 6.76×10^-16^). This was evidenced when levels of African ancestry were projected onto a map of North America (see **Fig. 5D**) revealing that westward samples (i.e., from the American midwest or California) have lower levels of African ancestry when compared to samples in the eastern seaboard at comparable latitudes. These results suggest that, in North America, the patterns seen under *k* = 8 emerge due to the different levels of African admixture (**Fig. 4D**, also **Fig. S11**).

**Figure 5:**
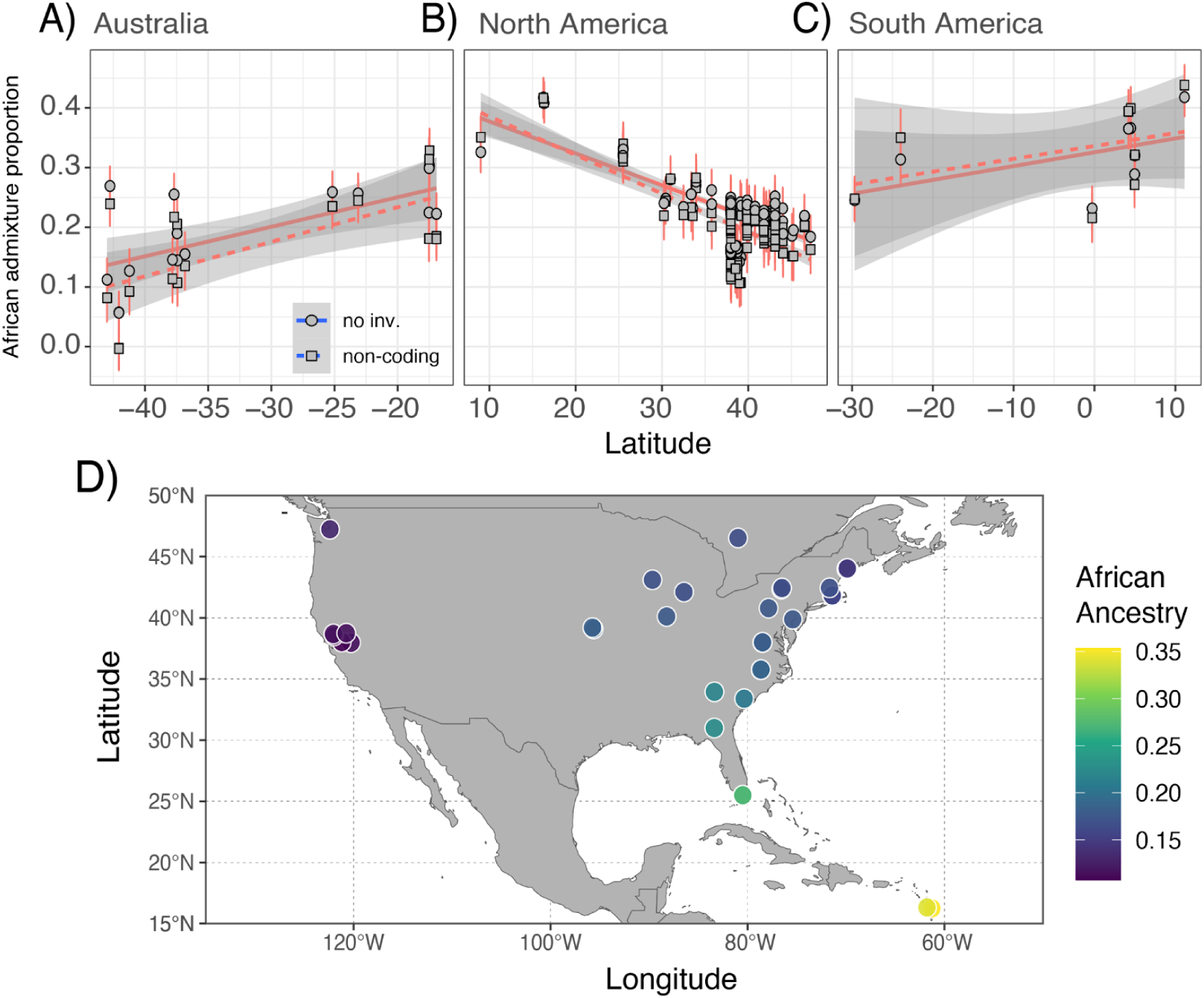
Patterns of admixture across the Americas and Australia. **(A)** Coefficients of linear admixture for Australia (excluding SNPs in inversions). **(B)** Same as A but for North America. **(C)** Same as A but for South America. **(D)** Map projection of levels of African ancestry in North American samples (note that the collapsed samples of Fournier-Level et al. 2019 were removed).

We further explored patterns of admixture using a two-pronged approach. First, we calculated the *f*_3_ statistic (Patterson et al. 2012; Gautier et al. 2022) using samples from North and South America as the targets of admixture and Europe and Africa as the “ancestral” populations. For African populations, we included samples from Cameroon, Egypt, Ethiopia, Morocco, Rwanda, South Africa, and Zambia. In total, we conducted 1,478,000 three-population comparisons (**Dataset S2**). Overall, all American populations displayed significant *f*_3_ tests (i.e., had a *Z*-score < -1.65), which confirms pervasive admixture (**Table S6;** also **Fig. S11**); these results do not appear to be driven by differences in read depth (*r*_signif_ *_f_*_3-RD_ = -0.6, *P* = 0.10) or by the number of flies included in the pool or synthetic pool (*r*_signif_ *_f_*_3-Nflies_ = 0.2, *P* = 0.40).

Lastly, we conducted a survey of genetic differentiation across the demographic clusters (see Materials and Methods: *Estimation of nucleotide diversity*). The overall differentiation was *F_ST_*= 0.050 ± 0.001 for autosomes and nearly twice as high for the X chromosome (0.091 ± 0.004; **Fig. 6A, orange**). These results were robust to the removal of heterochromatin regions and low frequency alleles (MAF < 0.05; **Fig. S12**). To quantify the level of differentiation between population groups defined by their continental cluster (**Fig. 4A**), we further relied on a hierarchical *F_ST_* model (Nei 1973), which consists of decomposing the total differentiation into an across-group (*F*_GT_) and a within-group (i.e., a composite label of continent and cluster; *F*_SG_) contributions, using unbiased estimators developed for Pool-Seq data (Gautier et al., *in prep.*). Note that here we refer to the overall differentiation under the hierarchical model as *hF_ST_* (with (1 - *hF_ST_*) = (1 - *F*_SG_)(1 - *F*_GT_)) to distinguish it from the standard *F_ST_* defined under a model without population groups (see above). As shown in **Fig. 6A**, *F_SG_* was always lower than *F_GT_*, demonstrating that there is less differentiation within than between most clusters. We evaluated the level of differentiation across all cluster-continent pairs by computing pairwise *F_GT_* (i.e., for each pair of regions the underlying populations were analyzed under a hierarchical *F_ST_* model with two groups), as shown on **Fig. 6B** (see results for *k* = 8 in **Fig. S13**). In general, all clusters involving Africa were consistently more differentiated than non-African groups. The highest level of differentiation was observed between Africa and EU-E (*F_GT_* = 0.22; **Fig. 6B**). Despite being located geographically between EU-W and EU-E, samples from the overlapping zone in Europe and Asia were more similar to EU-W than to EU-E (**Fig. 6B**). All populations in the Americas and Australia (i.e., “recent-expansion” populations) were more similar to each other than to Africa or Europe, reflecting a history of recent expansion and admixture between these two demes. Finally, we estimated the differentiation (i.e., standard *F_ST_*) within each cluster-continent level (**Fig. 6C**). Europe (cluster 2*_k_*_= 4_) exhibited the lowest levels of differentiation, and South America (cluster 4*_k_* _= 4_) the highest, which was essentially driven by a Brazilian and an Ecuadorian sample, the latter being separated in clustering at *k* = 8 (**Figs. 4B-D**).

**Figure 6:**
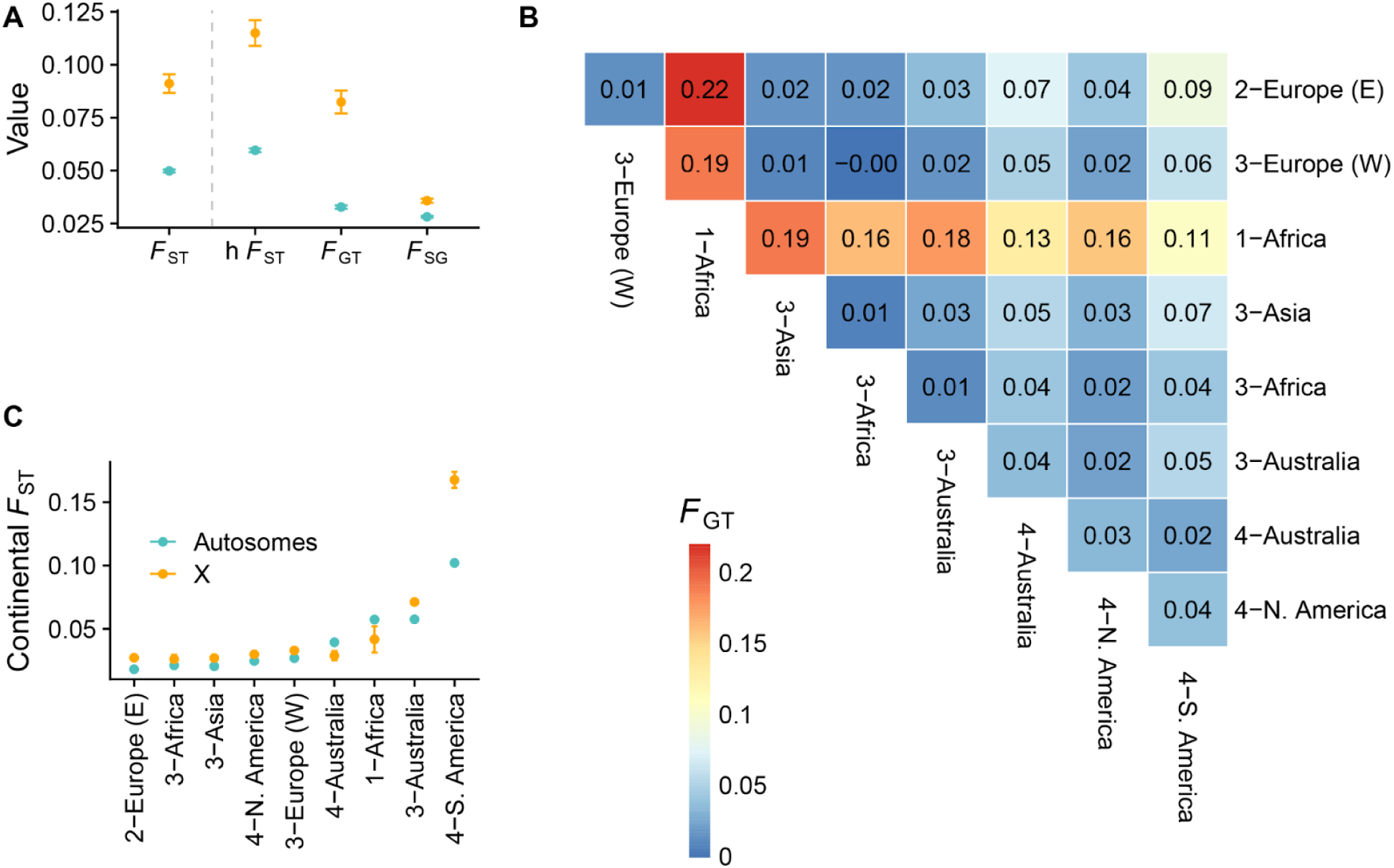
Genetic differentiation. **(A)** Values of the *F*_ST_ estimates over all DEST samples and their 95% CI (corresponding to ±1.96 s.e. estimated using block−jackknife with blocks of 50,000 consecutive SNPs). Note that the h*F*_ST_, *F*_GT_ and *F*_SG_ statistics were estimated using the hierarchical *F*_ST_ model, over all DEST samples grouped according to the *k* = 4 clustering analysis and their 95% CI. Colors indicate autosomes (blue) and X chromosomes (orange). **(B)** Pairwise comparisons between cluster-continents (under *k* = 4) results in a heatmap. In this plot, “1-Africa” refers to Sub-Saharan African populations, “3-Africa” refers to North Africa. The clusters “Australia-3” and “Australia-4” represent samples with low and high levels of African admixture, respectively. **(C)** *F*_ST_ estimates within clusters from the *k* = 4 analysis.

### Updated geographically informative markers improve predictive resolution of samples

Our previous release of DEST generated a panel of geographically informative markers (GIMs). The second release of our data gives us the unique opportunity to test the accuracy of our previously published markers. To this end, we applied our previously DEST 1.0 GIMs to our new data and we assessed the distance (*d*_hav_; as great circle distance, see *Materials and Methods*) between the predicted locality and the “real” locality as recorded in the metadata. Overall, both DEST 1.0 models trained at the level of “city” and “region” (i.e., resolution at the level of state or province), perform similarly well on the new data (*r* = 0.995, *P* = 2.2×10^-16^; **Fig. 7A**). Next, we aggregated the *d*_hav_ estimates at the level of continents (here we report only the results of the region model). We did this to assess whether the quality of our predictions vary as a function of continent. Overall, the best performance was observed in European samples (median resolution of ∼409 km to real location; **Fig. 7B**), followed by the North American samples, with a resolution of 794 km. Unsurprisingly, the worst predictions from the DEST 1.0 markers occurred when deployed on samples from South America and Australia, two locations that were not included in the first release (**Fig. 7B**).

**Figure 7:**
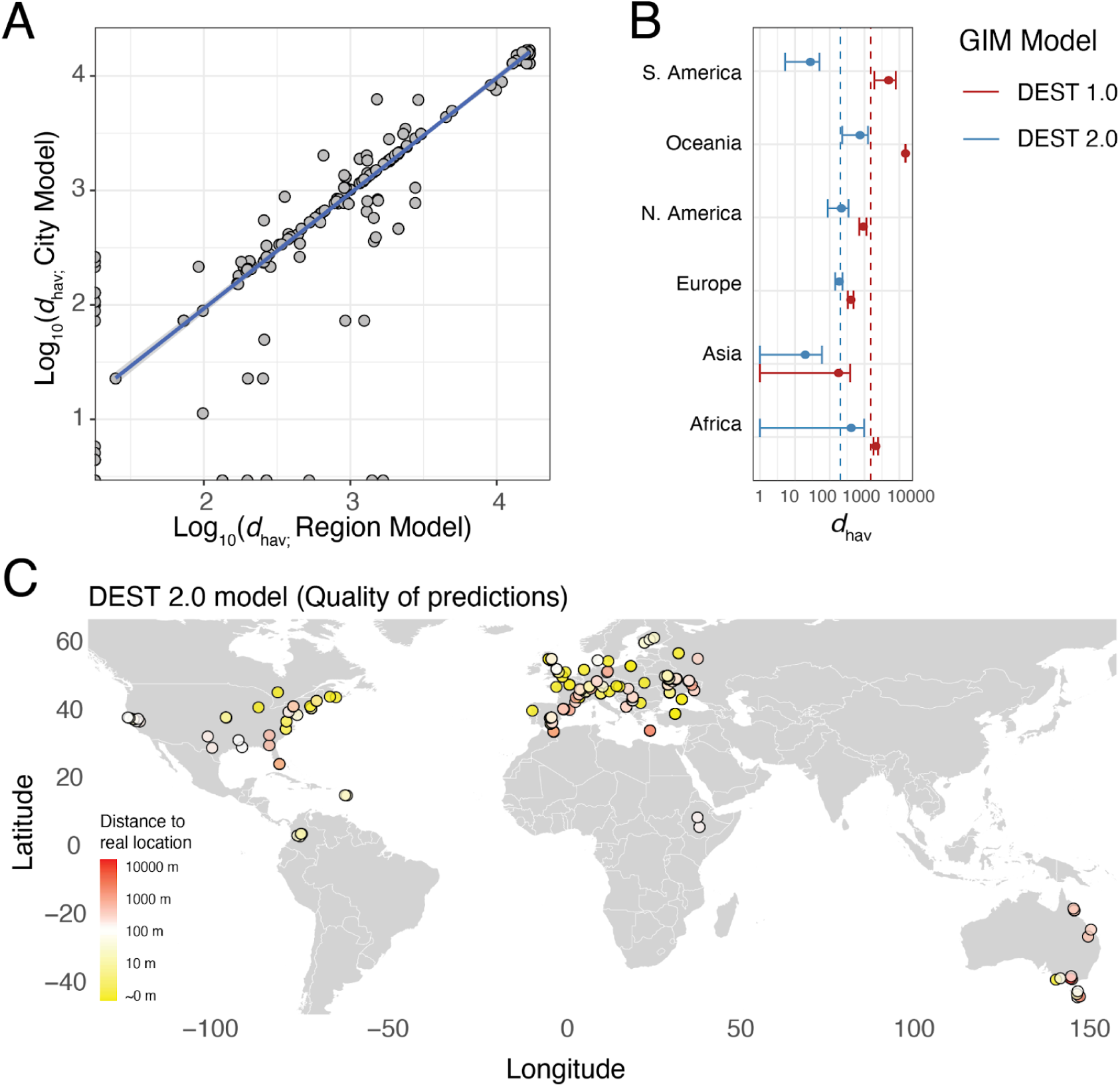
Geographically informative markers. **(A)** Bi-plot of *d*_hav_ from the 1.0 GIMs. City model (y-axis) and Region model (x-axis). (**B)** Mean and 95% confidence intervals (CIs) of *d*_hav_ for the 1.0 GIM and 2.0 GIM model (to improve readability the x-axis has been log_10_ transformed and CIs < 0 were set to 1; as 0 is logarithmically undefined). The mean distance to the true value is shown by dashed vertical lines (red for DEST 1.0, blue for DEST 2.0, models). (**C)** Quality of predictions for the GIM DEST 2.0 model. The color indicates the average distance between the real *d*_hav_ of a sample and its predicted *d*_hav_. Yellow are good predictions (accuracy = 0-10 m), white are “adequate” predictions (10-100 m), and red are poor predictions (1000-10000 m).

While our published markers performed well on samples from regions present in DEST 1.0, the addition of new regions to DEST required the generation of new GIMs. As such, we trained a new demographic model (DEST-GIM 2.0) including the new samples reported in this paper. Our new model was trained using the same workflow as DEST-GIM 1.0 (i.e., by retaining 40 PCs). Yet, the models differ in that DEST-GIM 2.0 was created by exclusively using non-coding SNPs as well as loci outside genomic regions spanning major cosmopolitan inversions. This new panel of GIMs is composed of 29,952 SNPs across all autosomes. Performance assessment of the new model by the *d*_hav_ analysis shows that DEST-GIM 2.0 performs similarly to the 1.0 version for existing locales (e.g., Europe or North America; **Fig. 7B**), yet they provide improved prediction accuracy for new regions (**Fig. 7B** and **7C**).

### Winter severity drives year-to-year levels of genetic variation in overwintering populations

While much of demographic research in *D. melanogaster* has focused on spatial patterns of genetic variation, there is strong evidence that temporal demography, driven by yearly cycles of summer “booms” and winter “busts”, can have strong and quantifiable effects on the frequency and levels of standing genetic variation in wild populations (Bergland et al. 2014; Nunez et al. 2024). For example, levels of post-overwintering (i.e., year-to-year) *F*_ST_ are generally higher than *F*_ST_ between samples collected within a growing season even though overwintering *F*_ST_ captures a smaller number of generations (1-2 generations) than comparisons within a growing season (ca. 10 generations). This observation has led to the hypothesis that strong bottlenecks due to overwintering alter the genetic composition of fly populations, both due to changes in the amount of genetic drift (Nunez et al. 2024) and due to seasonally varying selection (Bergland et al. 2014; Machado et al. 2021; Behrman and Schmidt 2022; Johnson et al. 2023). A prediction of this hypothesis is that the strength and intensity of winter, an ecological driver of yearly population busts, should be correlated with the levels of overwintering *F*_ST_ from one year to the next. To test this prediction, we investigated patterns of temporal structure in worldwide DEST samples and asked whether latitude (a proxy for winter severity) is correlated with the levels of year-to-year *F*_ST_.

For a given site, we assessed levels of *F*_ST_ between samples collected in two consecutive years (i.e., growing seasons) from the same locality. We implemented this analysis across 43 localities and estimated the relationship between mean year-to-year *F*_ST_ and latitude. We tested the hypothesis that higher-latitude populations with stronger winter conditions exhibit higher levels of year-to-year *F*_ST_. Indeed, we found a significant positive correlation between overwintering *F*_ST_ and latitude, yet the correlation is not monotonic. Using “broken-stick” regression (Muggeo 2003), we identified a change in the latitude-*F*_ST_ relationship at 50.3°N (**Fig. 8A** and **8E**). Samples below 50.3°N tend to have lower values of year-to-year *F*_ST_ as compared to those above 50.3°N (**Fig. 8B**) and the magnitude of correlation between latitude and *F*_ST_ varies before and after this latitude mark (**Fig. 8B**; *r*_all_ = 0.182, *r*_>50 lat_ = 0.333, *r*_<50 lat_ = 0.117; all *P* = 2.2×10^-16^). These correlations are statistically significant and outperform 500 random permutations where latitude is shuffled.

**Figure 8:**
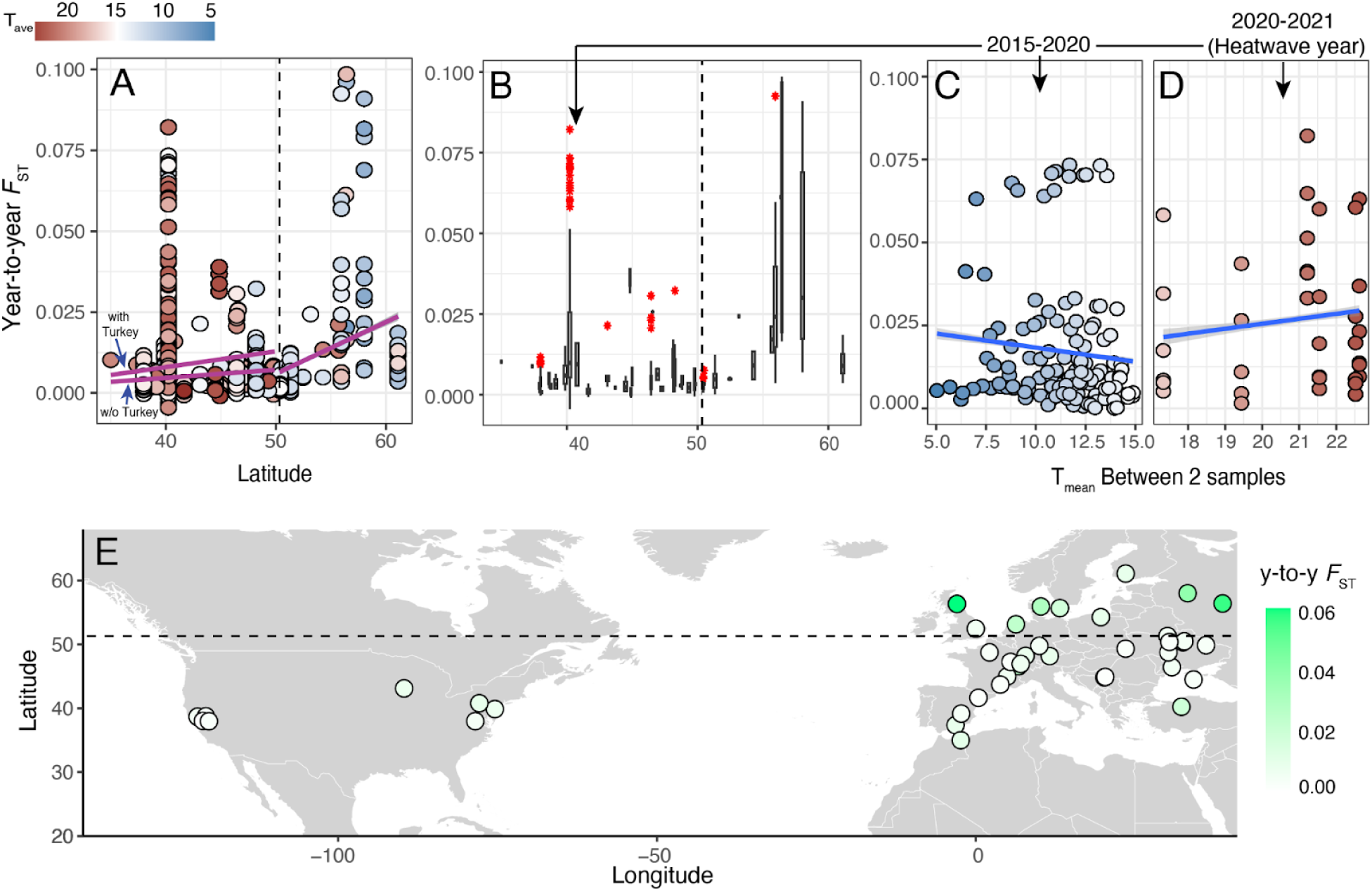
Temporal genetic differentiation due to overwintering. **(A)** *F*_ST_ values across DEST 2.0 samples as a function of latitude. Broken-stick regression and breakpoint is shown, for samples below latitude 50.3 the regression is shown with and without Turkey. The color indicates the mean temperature in Celsius between the samples for which the *F*_ST_ was calculated. **(B)** Distribution of year-to-year *F*_ST_ values across DEST 2.0 samples as a function of latitude, for comparisons spanning one winter only. Outliers (i.e., data above the 75th percentile) are shown in red. **(C)** Distribution of temporal *F*_ST_ values as a function of the mean temperature in Turkey (Yesiloz) samples for samples between 2015 and 2020 (logit transformed; correlation between *F*_ST_ and mean temperature; *r* =0.135; *P* = 4.60×10^-7^). **(D)** Same as B but for comparisons of 2020 and 2021, a historical heatwave year in Turkey and in southern Europe (correlation between *F*_ST_ and mean temperature; *r* = -0.100; *P* = 7.74×10^-13^). **(E)** Mean year-to-year *F*_ST_ overlaid over a world map of northern seasonal habitats.

A second finding of our year-to-year *F*_ST_ analysis was the discovery that several samples collected from Yesiloz, Turkey are outliers (red dots in **Fig. 8B**) among samples below the 50.3 latitude mark (see **Fig. 8A-B**). This pattern was most apparent when considering samples between 2020 and 2021 (**Fig. 8D**) relative to comparisons at other years (**Fig. 8C**) This signal in Turkey appears to be associated with a historical heatwave and unusually warm winters in 2021 (see discussion; **Fig. 8D**).

### Footprints of spatial adaptation to insecticides in Europe

The broad sampling inherent to DEST allows us to test hypotheses about spatial adaptation in wild flies. We first took a heuristic approach where we extracted all regions of the genome with high across-cluster differentiation (i.e., *F_GT_* > 0.2; see Results: *Population admixture and…*) and performed a gene ontology enrichment analysis of genes located in these regions of high differentiation (Kofler and Schlötterer 2012). Overall, we found an enrichment of genes associated with environmental adaptation such as responses to oxidative stress, metal ion and pesticides (**Table S7**). One of the strongest signals of population differentiation was observed for the region surrounding the gene *Cyp6g1,* a cytochrome P450 (Cyp) gene (**Fig. S14**; a result also observed in DEST 1.0), a well-known gene involved in resistance to DDT and neonicotinoid insecticides (Le Goff and Hilliou 2017). This signal was particularly high when comparing North America and European samples. Elevated *F*_GT_ was also observed when comparing South American and North American samples, but not when comparing South American and European samples (**Fig. S14**). These signatures of differentiation suggest different adaptations likely driven by distinct environmental pressures and insecticide exposure levels in each continent. To formally detect footprints of adaptive differentiation in our dataset we applied the “*Bayesian Population Association Analysis*” framework, *BayPass* (Gautier 2015; Olazcuaga et al. 2022) to DEST samples from European localities (irrespective of sampling year or season; 138 samples in total; **Fig. 9A**) and relied on the estimated *XtX*\* statistic to identify overly differentiated SNPs. The analysis identified two regions in chromosome 2R as candidates of local adaptation (12,188,558-12,126,181 and 14,826,182-14,976,108; **Fig. 9D**). Both these regions harbor several *Cyp* genes. For example, the window at ∼12 Mb contains *Cyp6g2*, and *Cyp6t3*, whereas the window at ∼14 Mb contains *Cyp6a22, Cyp6a19, Cyp6a9, Cyp6a20, Cyp6a21, Cyp6a8, and Cyp317a1*. These genes are associated with hormonal metabolism as well as responses to insecticides (Danielson et al. 1995; Le Goff and Hilliou 2017). We performed gene ontology enrichment analysis of genes within all *XtX*\* outlier regions and found an enrichment of terms such as “oxidation-reduction process”, “cellular response to radiation”, and “amide biosynthetic process”, reflecting results from *F_GT_* outlier regions above (**Table S8**).

**Figure 9:**
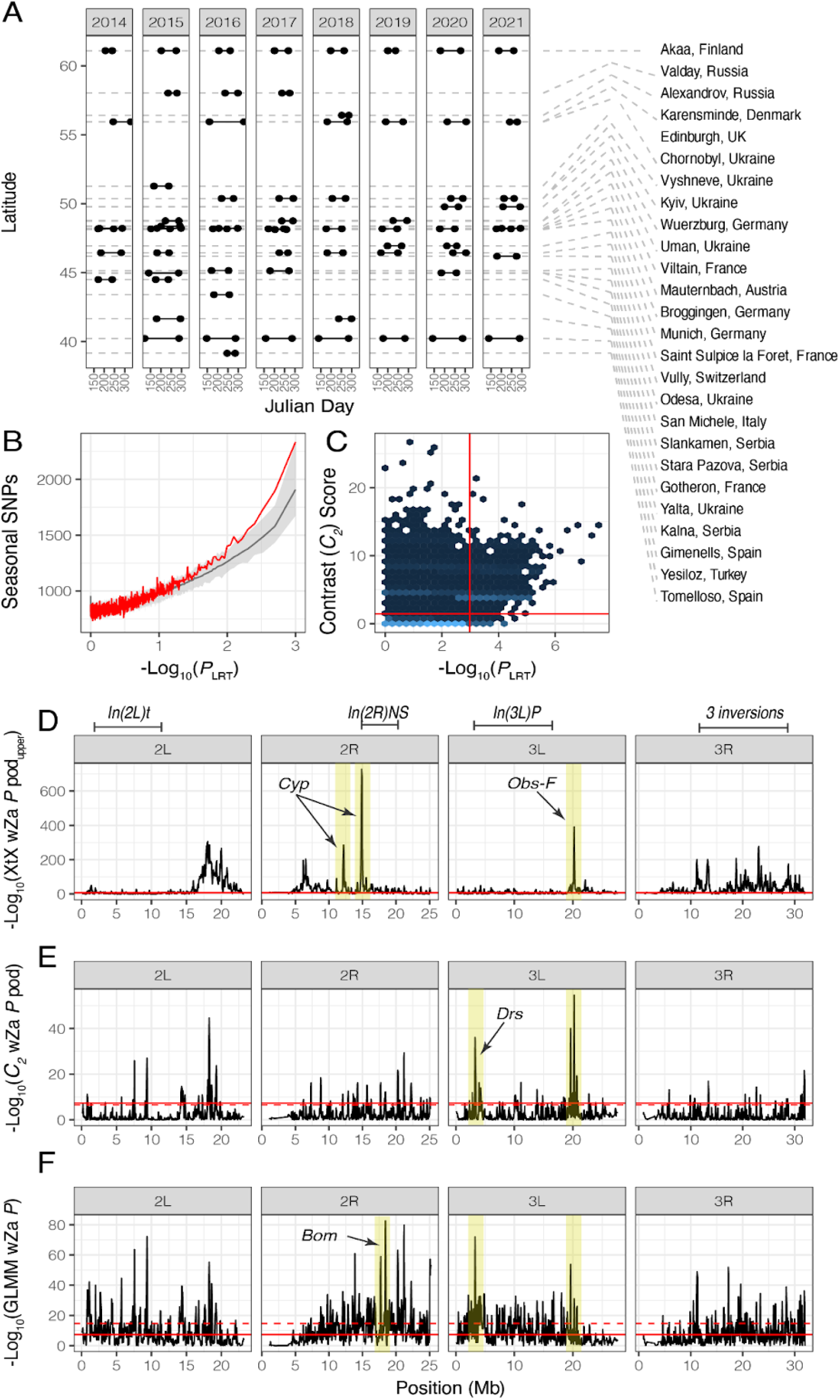
Local and seasonal adaptation in *Drosophila*. **(A)** Schematic of sampling for the seasonal analysis. In total, we used 138 samples collected in 26 European localities across an 8 year period. We selected localities where there were more than one sample per year and designated the first sample as “spring” and last sample as “fall”. There is no overlap between the samples used here and the samples used in seasonal analysis in Machado et al. (2020), Bergland et al. (2014), and Nunez et al. (2023). **(B)** GLMM seasonal adaptation scan. The plot shows the log_10_ transformed wZa *P*-value of the LRT of base and seasonal models. For A, B, and C, regions of interest are highlighted in yellow. Inversions are demarcated along the top of the figure. **(C)** We performed the contrast analysis using *BayPass* 2.4. The contrast score (*C*_2_ statistic) is the test statistic for the seasonal term, and follows a *χ*^2^ distribution with 1 degree of freedom. The x-axis is the -log_10_(*P*-value) from the GLMM. The red horizontal line represents the 99.9% significance threshold from the pseudo-observed data (POD) for ∼10M simulated sites. The red vertical line represents the 99.9% significance threshold from the permutations of the GLM analysis. **(D)** Bayesian local adaptation scan. The plot shows the log_10_ transformed wZa *P*-value of the local adaptation (*XtX*\*) *BayPass* analysis. **(E)** Bayesian seasonal adaptation scan. The plot shows the log_10_ transformed wZa *P*-value of the contrast (*C*_2_) adaptation *BayPass* analysis. **(F)** Results of the GLMM analysis. The permutations are shown in gray (95% confidence intervals) and the real data in red. There are more SNPs with low seasonal p-values than expected by permutations.

### Antimicrobial peptides are enriched among continent-wide targets of seasonal adaptation

We explored signals of seasonal evolution in DEST using paired spring-fall collections from Europe. In order to ensure that this test was not influenced by signals from previously analyzed data, we only used samples that were not included in previously published analyses (i.e., Bergland et al. 2014; Machado et al. 2021; Nunez et al. 2024; **Fig. 9A**). First, we ran the *BayPass* model including both the Ω matrix as a demographic prior as well as categorical “spring” or “fall” labels (defined by the first and last sample collected in a locality within a year) in a contrast analysis. Under these conditions, *BayPass* outputs the *C*_2_ statistic that quantifies the degree of association of allele frequency with season. We identified significant *C*_2_ values using a simulation approach that is part of the *BayPass* workflow (see Materials and Methods: S*cans for adaptive differentiation*; **Dataset S3**). We observe that several regions across the *Drosophila* genome are enriched for signals of parallel seasonal evolution (**Figs. 9D E, F**). A notable example appears in chromosome 3L (3,222,669-3,422,464), inside the region spanned by the inversion *In(3L)P*, where we observe the antimicrobial peptide *Drosomycin* (*Drs*) as well as several *Drs*-associated genes (i.e., *Drsl2*, *Drsl3*, *Drsl4*, *Drsl5, Drsl6*). In view of previous observations of seasonal allele frequency oscillations in several immune genes, this result suggests functional shifts in immune tolerance and resistance across seasons in natural populations (Behrman et al. 2018). We performed gene-ontology enrichment analysis of all genes within *C*_2_ outlier regions (**Table S9**). We found an enrichment of, among other terms, genes associated with “alcohol dehydrogenase (NAD) activity”, including the gene *Adh* itself (**Table S10**).

We conducted an enrichment analysis comparing our *C*_2_ SNPs (in the top 0.0001 %) with loci reported in previous seasonal studies, done mostly in North American populations (i.e., FDR < 0.3 in Bergland et al. 2014; Top 1% SNPs in Machado et al. 2021), to assess whether seasonal SNPs in Europe are also likely to be seasonal in North America. Our results indicate no significant enrichment of North American seasonal SNPs among our European *C*_2_ SNPs (**Fig. S15**). Indeed, when compared to Pennsylvania data from Bergland et al. (2014), we observed a significant deficiency of these targets at both a global level (*P* = 0.024; **Fig. S15A**) and specifically on chromosome 3L (*P* = 0.0055).

Beyond the *C*_2_ analysis, we implemented a generalized linear mixed model (GLMM) using the spring/fall seasonal labels, showing a global enrichment of seasonal SNPs relative to permutations (**Fig. 9B**). Comparing GLMM and *BayPass* results, we found a large number of SNPs exceeding the simulated 99.9% significance threshold for the *C*_2_ statistic (**Fig. 9C**, red vertical line), with the *C*_2_ and GLMM models producing a similar set of candidate SNPs (**Fig. 9C**, red horizontal line). Likewise, a sliding window wZa analysis (Booker et al. 2024) of the GLMM results (window size of 100 kb, step size of 50 kb) identified the *Drs* region as a hotspot of seasonal adaptation (as in the *C*_2_ analysis), and also revealed a second region of interest on chromosome 2R (18,376,129-18,475,992). This region contains several *Bomanin* genes (abbr. *Bom*; e.g., *BomBc1, BomT1, BomS1, BomBc2, BomS6*) known to play key roles in *Drosophila* antifungal responses (Xu et al. 2023). A region on 3L, near 20,172,964-20,271,926 bp, notable for harboring adjacent signal peaks across analyses of seasonal and local adaptation (see **Figs. 9D, 9E, 9F**; yellow band), contains *obstructor-F* (*obst-F*), a gene previously reported as a candidate of insecticide adaptation (Campo et al. 2013; Bogaerts-Márquez et al. 2020).

## Discussion

### A unified resource for wild *Drosophila* genomics

*D. melanogaster* is a cosmopolitan species with resident populations across all human-inhabited continents that evolves adaptively in response to spatially-varying and temporally-fluctuating selection in semi-natural settings and the wild (clinal patterns reviewed in Adrion et al. 2015; seasonal patterns reviewed in Johnson et al. 2023). To achieve a comprehensive understanding of the evolutionary patterns within this species, we need to create panels of variation sampled across wide geographical scales and densely across time. This is not a trivial undertaking for any single lab to achieve. The original impetus behind DEST was to generate a unified dataset and workflow that would capitalize on the collaborative efforts of labs and consortia around the world (Kapun et al. 2021). DEST 2.0 expands data on the original release by adding twice as many new samples as the original release.

Overall, the incorporation of the aforementioned data into the dataset showcases the flexibility and capacity for growth of DEST, as a centralized and well annotated repository of *Drosophila* genomics. Furthermore, the DEST 2.0 *Dockerized* pipeline now allows for pools generated using single-end sequencing approaches to be incorporated into its workflow, hence allowing for older pooled data sets to be included in DEST analyses. We plan to continue maintaining and updating the DEST workflow, with potential future expansions to explore other *Drosophila* species and additional data types. To keep pace with the influx of new genomic data, we have upgraded the DEST genome browser to the latest version of JBrowse, which has better scalability and performance when displaying large datasets (Diesh et al. 2023).

### Heterogeneous patterns of recombination in DEST samples

This release also includes genome-wide recombination rate estimations for 75 representative populations. In comparison to the findings of previous studies (Comeron et al. 2012; Adrion et al. 2020) our own estimates show a reduction of approximately threefold. This discrepancy may be attributed to the combination of our methodological approach and the nature of our data. The deep learning approach of *ReLERNN* (Adrion et al. 2020) is dependent on allele frequencies, and it is thus possible that levels of genetic polymorphism may affect the estimation of levels of recombination rate. In our analyses, we estimated allele frequencies on SNPs that were called with very conservative and stringent filtering methods. Furthermore, the polymorphism data were obtained from Pool-seq data from derived European and North American populations, which exhibit lower levels of genetic polymorphism (approximately two- to three-fold; e.g., Ometto et al. 2005) than the ancestral African populations used in Adrion et al. (2020). Accordingly, there is a strong, and significant, correlation between the number of SNPs and the average recombination across the 75 populations (Spearman’s rho = 0.835, S = 11624, *P* < 1.0×10^-25^; R^2^ = 0.672). It is thus possible that our estimations can be approximated as a population-scaled effective recombination rate (*ρ*) rather than the actual crossing-over rate (*r*; where *ρ* = 4*N*_e_*r*). A comparable finding was observed in the case of wild barley (Dreissig et al. 2019). It seems also probable, however, that our populations can indeed be characterized by heterogeneous levels of recombination, as has been reported by numerous studies in *Drosophila* (e.g., Hunter et al. 2016; Samuk et al. 2020; Wang et al. 2023).

### New insights into ancestral and recent fly phylogeography

The prior releases of DEST and similar datasets (Kapun et al. 2020; Kapun et al. 2021; Machado et al. 2021) characterized fine-grained levels of population structure within Europe, and dated their divergence at around ∼1,000 ya. In this paper, we expanded the repertoire of samples available for demographic inference and phylogeographic analysis.

In the Americas and Australia, our data recapitulate published patterns of African admixture in North American fly populations (Kao et al. 2015; Bergland et al. 2016; Corbett-Detig and Nielsen 2017). Notably, in South America and Australia, while not significant, our results show a reversed trend with latitude, relative to North America (**Fig. 5A-C**). These support the general hypothesis of higher African admixture in equatorial populations relative to poleward ones, consistent with two separate introductions of *D. melanogaster* to the Americas. It is likely that the African ancestors entered the Americas through the Caribbean. In this region, the earliest record of *D. melanogaster* occurred in Cuba in 1862 (Sturtevant 1921), and it was first documented in Florida in 1894 (Keller 2007). While it is always important to consider that species distributions data may be incomplete, the entomological surveys conducted in the USA during the 1880s are extensive and they do not mention earlier records of the species under any of its old taxonomic names (i.e., *D. ampelophila or D. uvarum*; see Keller 2007). The origin and timing of European immigration is more complex. Notably, European entomological surveys only describe the presence of *D. melanogaster* as a “common” species in Central Europe (Sturtevant 1921), with reported sightings in German cities like Kiel or in Austrian towns in the 1830s (Keller 2007). Consistent with this chronology, the first recorded samples in North America come from New York in 1875 (Lintner 1882; Keller 2007). Thus, while African flies may have been in the Americas since the 1860s, it is possible that the African-European admixture cline in USA’s eastern seaboard originated later, during the late 1880s.

In Europe, the overlap zone we observed inside the continent (in the *k* = 8 analysis) is notable since its placement closely mirrors the “suture zones” (Remington 1968) of other species such as *Bombina* toads (Hofman et al. 2007)*, Leuciscus cephalus* (Hewitt 2011), and *Mus musculus* (Ďureje et al. 2012). In our analyses, we tested whether this overlap zone is a zone of admixture between EU-E and EU-W. We reject this model and suggest that the overlap zone is a subpopulation of EU-W. These results are puzzling, and echo findings from our previous release (Kapun et al. 2021), whereby the levels of gene flow in this area appear to be asymmetric in favor of EU-W (e.g., as reported by Kapun et al. 2021, EU-W→EU-E as 0.209 flies/gen vs. EU-E→EU-W as 0.178 flies/gen). These findings are supported by our supplementary *F*_ST_ analyses that include the overlap zone (e.g.; *F*_ST_ [EU-W vs. Overlap] = 0.00; *F*_ST_ [EU-E vs. Overlap] = 0.01). As it stands, these patterns may indicate the action of a non-neutral force confounded with the complex demographic history of *D. melanogaster* in Europe, to be explored in future work.

### Inferring targets of adaptation across time and space

The complex patterns of spatial population structure that we have described above are likely to alter the adaptive capacity of fly populations. Indeed, a recent genomic analysis of the sibling species *D. simulans* across continents revealed that demographic ancestry, and not shared selection regime, is a better predictor for the genetic basis of local adaptation to thermal stressors (Otte et al. 2021). These results highlight that assessing footprints of adaptation requires robust controls for the complex demographic structure of species. We implemented the *BayPass* framework (Gautier 2015; Olazcuaga et al. 2022) to discover targets of spatially and temporally fluctuating selection across Europe. This framework is flexible, as it incorporates priors from population structure (via the Ω matrix) and, optionally, environmental variables (either as factors or covariates).

Our analyses of spatial adaptation reveal signatures of continent-wide differentiation around cytochrome P450 genes (e.g., *Cyp* genes) in 2R (**Fig. 9**). Follow-up analyses using estimates of across-group differentiation (*F*_GT_) revealed that these genes are highly differentiated in comparisons between North American populations vs. both European and South American populations (**Fig. S14**). Given that *Cyp* genes are important players in insect detoxification pathways and have been implicated in the evolution of insecticide resistance (Le Goff and Hilliou 2017), these findings suggest that flies have experienced continent-wide adaptation to different histories of land and pesticide use. While further experimental validation is needed to disentangle the particular gene targets and drivers of selection, these data highlight the power of DEST to reveal the genetic bases of local adaptation to paralleled stressors.

We also explored patterns of temporal divergence in response to seasonality. Previous work has shown that seasonal adaptation, via adaptive tracking (Botero et al. 2015), is a ubiquitous, and important, evolutionary force affecting patterns of genetic variation across the genome of *Drosophila* (Bergland et al. 2014; Kapun et al. 2016; Machado et al. 2021; Rudman et al. 2022; Bitter et al. 2024; Nunez et al. 2024). Here, we used the DEST 2.0 data to revisit footprints of seasonal adaptation across samples not used in previous analyses. Using this dataset, we tested the hypothesis that seasonal adaptive tracking is a general phenomenon of worldwide temperate *Drosophila*. One challenge associated with testing this hypothesis is determining the appropriate covariate (e.g., temperature, humidity, rainfall) and the timeframe of selection (e.g., 0-15, 0-30 days prior to collection) to use in the model. For example, Nunez et al. (2024) showed that, in Virginia, the best seasonal model used the temperature 0-15 days prior to collection as a covariate. Yet, in Europe, Humidity 0-30 and 0-60 prior to collection days were the best models for EU-E and EU-W respectively. Therefore, we used a contrast framework using the seasonal labels (i.e., “spring” and “fall”) as comparison factors. This approach had been successfully used in the past by Bergland et al. (2014) and Machado et al. (2021) and allowed us to surmount the challenge of covariate selection.

We implemented a test of seasonality in a two-pronged approach using both the *BayPass* and the GLMM framework. Our results show multiple regions of interest across the genome that are concordant across both BayPass and GLMM. For example, it highlights a region on 3L that encodes for *Drosomycin* and *Drosomycin-like* genes (**Fig. 9D**), canonical antifungal defense loci (Zhang and Zhu 2009), as a continent-wide hotspot of seasonal adaptation (**Figs. 9C, 9F**). These findings are noteworthy, as fungal communities are known to vary drastically across seasons driven by changes in soil moisture, temperature, and carbon availability (Schadt et al. 2003). Furthermore, the analysis also reveals a region of interest on chromosome 2R containing *Bomanin* genes that are also associated with antifungal defense (Xu et al. 2023). Another gene of interest is *Obstructor-F, a* gene that has several functions and that has been associated with pesticide response (Campo et al. 2013).

Our gene-ontology enrichment analysis for targets of seasonality highlighted “alcohol dehydrogenase activity” —including the gene *Adh* itself— as being enriched among outlier regions. This is significant because patterns of genetic variation in *Adh* have long been recognized as classical examples of ecological adaptation (Kreitman 1983; Berry and Kreitman 1993). However, recent discussions have emphasized that the specific agents of selection acting on this gene remain unclear, with some suggesting temperature-driven balancing selection (Siddiq and Thornton 2019). We also assessed whether the seasonal SNPs observed in our *C*_2_ analysis from Europe are enriched in seasonal datasets generated mostly from North American populations (Bergland et al. 2014; Machado et al. 2021). Our results showed no enrichment (or under-enrichment; see **Fig. S22**) between the datasets compared. In other words, these results suggest that the genetic basis of seasonality is different between continents. This finding is consistent with previous studies positing that population ancestry is a more important predictor of adaptive genetic architecture than the existence of paralleled selection regimes (Otte et al. 2021).

Overall, our seasonal analyses reveal three major takeaways. First, they reveal that seasonal adaptive tracking is a detectable phenomenon across the temperate range of *D. melanogaster*. Yet, they also suggest that adaptive tracking may be driven by both natural and anthropogenic stressors, and that the specific loci that drive adaptation may be strongly shaped by genetic ancestry. Second, the data highlight a large role of pathogen response genes as major players in worldwide seasonality (Behrman et al. 2018). These findings suggest that follow-up studies of seasonality should take a more comprehensive approach to incorporate both abiotic (e.g., temperature) and biotic (e.g., pathogen) views of “seasonality.” And third, our findings showcase an inherent strength of the *BayPass* model to successfully disentangle the dynamics of spatial and temporal adaptation in wild populations. Further expansions of the DEST dataset will facilitate more granular exploration of adaptive tracking driven by spatially and temporally fluctuating selection.

### The impacts of overwintering demography on genetic variation

The results highlighted above showcase the power of DEST to examine fine-grained patterns of evolutionary change occurring within each population. Yet, seasonal adaptive tracking is not the only process at play in temperate habitats. As the seasons change, *Drosophila* populations expand and contract depending on resource availability (Atkinson and Shorrocks 1977). Indeed, the establishment and range limits of many insect species are tied to their ability to survive winter (Lawton et al. 2022). Previous work has suggested that local fly populations grow to their largest possible size during the summer months (Atkinson and Shorrocks 1977; Sanchez-Refusta et al. 1990; Gleason et al. 2019; Bangerter 2021) and drastically decrease in size following the onset of winter, when resources are scarce and reproduction is suppressed, leading flies to diapause and overwinter until the next growing season. These seasonal demographic cycles, called “boom-and-bust” demography, can result in yearly bottlenecks of up to ∼97% in the “local” population (Nunez et al. 2024), and thus are likely to have fundamental consequences for standing genetic variation.

One important question related to these boom-and-bust dynamics is whether populations that experience different severities of winter (harsher vs. milder) show elevated levels of year-to-year differentiation. We explored this question using year-to-year *F*_ST_ and tested the hypothesis that populations with harsher winters have, on average, larger levels of year-to-year *F*_ST_. Our results support this hypothesis, revealing positive correlations between *F*_ST_ and latitude, particularly for samples collected at latitudes higher than 50.3°N (**Fig. 8A** and **8E**). These patterns suggest that habitats with colder, harsher winters typical of higher latitude habitats impose stronger bottlenecks on overwintering flies relative to lower latitude habitats. One notable exception to the pattern of year-to-year *F*_ST_ was found in the Turkish samples. There, populations in 2021 showed an unexpected positive correlation between *F*_ST_ and temperature (**Fig. 8D**; relative to patterns at previous years at the same site, **Fig. 8C**). These patterns may have arisen as a result of the harsh weather conditions of southern Europe in 2021. During that period, weather anomalies created unusually warm winters as well as the hottest and longest summer heat waves in the region’s recent history (Lhotka and Kyselý 2022). These extreme heat waves may have affected flies both directly, through physiological thermal challenges, and also indirectly by affecting their food sources.

Overall, our findings provide two major insights into the temporal structure of *D. melanogaster* populations. First, we showed that overwintering bottlenecks are associated with the severity of winter across habitats. Second, that there is a predictable relationship between the strength of winter and the genomic consequences of overwintering in fruit flies.

### Future directions

In conclusion, our findings not only highlight the power of DEST as a resource for fly biologists but also its promise and potential for growth. Indeed, as more temporal samples continue to be added, more detailed gene-environment association studies will undoubtedly shine a light on the drivers of selection across worldwide habitats. Our data may also be used in order to parameterize temporally and spatially explicit population genetic simulations which, combined with climate change forecasting datasets, will help to model rapid evolutionary responses under various climate scenarios. Lastly, as our consortium continues to grow, we are working to include a variety of other *Drosophila* species into DEST. Such multi-species data will be pivotal to assess the evolutionary dynamics of adaptive tracking across the phylogeny.

## Materials and Methods

### Sample mapping and SNP discovery using the DEST mapping pipeline

Samples were mapped to the *D. melanogaster* hologenome using the pipeline described in our first release (Kapun et al. 2021). This pipeline consists of a combination of genomic tools (fast-qc [v0.12.1], Cutadapt [v2.3] (Martin 2011), BBMap [v38.80] (Bushnell et al. 2017), BWA-mem [v0.7.15] (Li 2013), Picard [v3.1.1], SAMtools [v1.9] (Li et al. 2009)) in a Docker container. For our current release of DEST (2.0), we have updated the Docker container to enable mapping of reads sequenced in both paired-end (PE) and single-end (SE) configuration. This new version of the pipeline can be found in Dockerhub (https://hub.docker.com/) as destbio/dest_freeze2:latest. SNP calling was performed using the PoolSNP algorithm (Kapun et al. 2020). For SNP calling, we used the default parameters optimized in the first release of DEST (Kapun et al. 2021). The SNP calling step as well as genome annotation with SNPEff (v5.2; Cingolani et al. 2012) were automated using SnakeMake (Mölder et al. 2021). We provide ready to use outputs of the DEST pipeline both in variant call format (VCF) format as well as in genomic data structure (GDS) format (Zheng et al. 2012). The entire DEST pipeline can be found on GitHub at: https://github.com/DEST-bio/DESTv2.

### Previously published datasets added to DEST 2.0

We incorporated data from previously published studies (Reinhardt et al. 2014; Svetec et al. 2016; Fournier-Level et al. 2019; Lange et al. 2022; Nunez et al. 2024). These data were added to DEST by processing the raw sequences using the Docker pipeline. These new samples include: 37 samples from Nunez et al. (2024), 16 samples from Fournier-Level et al. (2019), two samples from Hoffmann et al. (2002), 17 samples from Lange et al. (2022), eight samples from Reinhardt et al. (2014), and one sample from Svetec et al. (2016). Comprehensive metadata for these samples is included in **Table S1.** Samples from Fournier-Level et al. (2019) consist of multiple replicates from the same locality each with low coverage. Accordingly, we collapsed all replicates from each site into a single “consolidated” library (see “Collapse” category; orange squares in **Fig. 1C**), each with read depths of ∼60X.

### Filtering parameters

We filtered SNPs and samples using metrics and tools described in our first release (Kapun et al. 2021). In brief, we 1) calculated the levels of contamination by congenerics, 2) levels of read duplication in the sequencing run, 3) proportion of SNPs with missing allele frequency data, 4) ratio of synonymous to non-synonymous polymorphism (*p*_N_/*p*_S_), 5) nominal coverage, and 6) the effective coverage. Levels of contamination by congenerics refers to the amount of non-*D. melanogaster* flies accidentally sequenced in pools.

We assessed contamination using a two-pronged approach. First, we assessed levels of competitive mapping of reads to the genomes of *D. melanogaster* (RefSeq: GCF_000001215.4) and *D. simulans* (RefSeq: GCF_016746395.2). *D. simulans* and *D. melanogaster* can be difficult to differentiate in the wild and the wrong species may be sequenced by accident. The specifics of competitive mapping are discussed in the methods of the first release (Kapun et al. 2021). Our second approach uses a *k*-mer counting method that can be directly applied to raw read files and is flexible for multiple species that are represented or closely related to those represented in the target *k*-mer dictionary. This approach is described in (Gautier 2023). Next, we generated in-silico pools consisting of mixtures of panels of inbred *D. melanogaster* (Mackay et al. 2012) and *D. simulans* (Signor et al. 2018). We generated these in-silico pools by varying the mixture levels of the two species. By analyzing these pools, we show that both the competitive mapping and the k-mer approach are accurate (**Fig. S3A**), with the competitive mapping approach slightly over-estimating contamination (by 2.3% max) and the *k*-mer approach slightly under-estimating contamination (by 6% max).

The levels of read duplication were extracted directly from the BAM files by mining the “mark_duplicates_report” output using a custom R script. Missing data was assessed by counting the number of sites reported as “NA” in a particular pool. The *p*_N_/*p*_S_ statistic was calculated using the SNP annotations derived from SNPEff using custom script (see GitHub). The nominal, genome wide, read depth (RD) is extracted directly from the BAM file using a custom script (see GitHub). Note that the per-site RD is a standard output of PoolSNP.

### Masked gSYNC files

Prior to SNP calling, we masked positions in each gSYNC file, which is a genome-wide extension of the SYNC file format (Kapun et al. 2021) for each sample based on minimum and maximum read depth thresholds, as well as on proximity to putative indel polymorphisms as identified by GATK IndelRealigner v3.8.1 (DePristo et al. 2011). In addition, we masked regions associated with repetitive elements identified as fragments of interrupted repeats by Repeat Masker (Smit et al. 1996; Jurka 2000), microsatellites and simple repeats identified by Tandem Repeat Finder (Benson 1999), repetitive windows identified by Window Masker and SDust (Morgulis et al. 2006), and transposable elements and other repetitive elements identified by Repeat Masker (all obtained from the UCSC Genome Browser), using the custom python script MaskSYNC_snape_complete.py as previously described in Kapun et al. (2021). Importantly, the position of these masked sites are stored in BED file format, which allows accounting for masked sites both in mono- and polymorphic positions when calculating unbiased site-specific averages for population genetic statistics as described below in the section “Estimation of nucleotide diversity” (see also Kapun et al. 2020).

### Effective read depth

In addition to the nominal RD, multiple downstream analyses in this paper use the “effective RD” metric (n_e_). This is a Pool-Seq specific metric that corresponds to the number of individually genotyped chromosomes, after accounting for the double binomial sampling that occurs in Pool-Seq (Kolaczkowski et al. 2011; Feder et al. 2012; Gautier et al., 2013). An estimate of n_e_ for a Pool-Seq sample can be defined as

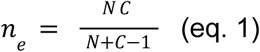

where N is the haploid sample size of the pool (i.e., number of pooled chromosomes) and c is the nominal RD at a given position or average across the genome (see **Text S1** for further details on the derivation of eq. 1 and for a more general formula applicable to collapsed Pool-Seq sample).

### Recombination landscape

We inferred the genome-wide recombination landscape for 75 of our samples using ReLERNN v1.0.0 (Adrion et al. 2020). The samples were selected to cover the entire spatial distribution of the DEST 2.0. sampling and based on the coverage sequencing depth (mean = 68.3, SD = 35.8, min. = 32, max. = 234), which was chosen to be as high as possible to maximize the reliability of the allele frequency used by ReLERNN to estimate recombination (**Table S1**). We used BCFtools (Danecek et al. 2021) to extract allele frequency of all biallelic SNPs with a frequency > 0.01 and read depth > 10. The resulting data was used to run ReLERNN. The parameters used in ReLERNN *simulate* module were as follow: assumed per-base mutation rate: --assumedMu 3.27×10^-9^; assumed generation time (in years): --gentime 0.08; and upper rho/theta ratio --upperRhoThetaRatio 10. For the train module, we applied a MAF of 0.01 (--maf). For the prediction module, we considered windows with a minimum number of 50 sites (--minsites). Following the developers’ recommendation, we let the program select the optimal size of the non-overlapping windows on which per-base recombination rates were predicted. To allow comparisons between samples, we estimated the average per-base recombination rates in larger 200 kb non-overlapping sliding windows by combining the raw rates estimated in each ReLERNN-selected window weighted by the fraction of the overlap with the corresponding 200 kb sliding window. Using the same approach, we also calculated the recombination landscape using the raw data of (Comeron et al. 2012), which are significantly correlated with our estimates for most of the populations (**Table S11**). Recombination rates are available in the genome browser.

### Estimation of nucleotide diversity

We conducted population genetic analyses using *npStat* (Ferretti et al. 2013). Out of the 530 high-quality samples, we used a subset of 504 samples for which we also had the masked bam files, which were necessary to compute the statistics. The remaining 26 samples do not have a masked bam file as they were incorporated from the DGN data. For those samples, diversity statistics come from DEST 1.0 data (Kapun et al. 2021). Standard nucleotide diversity statistics were first directly estimated from each *bam* file, for non-overlapping windows (10 kb, 50 kb or 100 kb) over the whole genome, using the estimators for Pool-Seq data developed by Ferretti et al. (2013). Only positions covered by at least two reads and less than 250 reads with a min quality > 20 were considered in the computations (*-mincov 2 -maxcov 250 -minqual 20* options) and windows with less than 9,000 remaining positions were discarded. We further calculated window-specific average estimates for each sample, using window sizes of 10k, 50k and 100k (i.e., window size that are displayed in the genome browser) using a custom Python script (BED2Window.py).

### Analyses of chromosomal inversions

Based on previously identified inversion-specific marker SNPs (Kapun et al. 2014), which are in tight linkage with the breakpoints of the common cosmopolitan inversions *In(2L)t*, *In(2R)NS*, *In(3L)P,* and *In(3R)Payne* and of the rare cosmopolitan inversions *In(3R)C, In(3R)K* and *In(3R)Mo*, we estimated sample-specific inversion frequencies based on the median of the frequencies of inversion-specific alleles across SNP markers for a given inversion following the approach in Kapun et al. (2014). To test for associations between inversion frequencies and geographic variables, we partitioned the data by continent and analyzed each inversion separately. We fit general linear models including arcsine square-root transformed inversion frequencies as dependent variables, which accounts for the skewed variance distribution in binomial data when normality is assumed. We included latitude, longitude and sampling year as independent variables and tested for the effect of the independent variables and all possible interactions with a likelihood ratio test. While we considered latitude and longitude as continuous numerical variables, we treated year as a categorical factor to account for the sparse sampling across years at most locations.

### Principal Component Analysis (PCA)

Global population structure analyses were done using the PCA algorithm implemented in the FactoMineR v2.4 package (Lê et al. 2008). For these analyses, we included all available samples that passed the filter in DEST 2.0. We include all biallelic SNPs in autosomes provided they had less than 1% missing data and a mean allele frequency greater than 1% (across all samples). We thinned the dataset by only selecting SNPs that were 500 bp apart from each other, reducing the dataset to 168,408 SNPs. Note that we ensured that this PCA was robust to variations in read coverage and haploid pool size by comparing the estimated PCs with those obtained with a random allele PCA, as implemented in *randomallele.pca*() from the R package *poolfstat* (v 2.3.0, Gautier et al., *in prep.*; **Fig. S7**).

### Demographic inference with *moments*

We fit demographic models to subsets of the DEST 2.0 variant data with the Python package *moments* (Jouganous et al. 2017). We adapted *moments* code to construct site frequency spectra (SFSs) from autosomal SNPs from the Pool-Seq VCF file, subset to include only the pool with greatest effective sample size (*n*_e_) from each locality in order to avoid geographic sampling bias. For simplicity, we normalized population-specific sample sizes to the average *n*_e_ of respective subsets of pools in consideration. For different subsets of the data, we constructed *demes*-type models (Gower et al. 2022) dubbed “one-population,” “split,” “two-splits,” and “admixture” (see **Fig. S9**) in order to infer demographic parameters of global *Drosophila* populations while simultaneously performing likelihood-based model selection. A significant limitation of SFS-based demographic inference (e.g. Gutenkunst et al. 2009; Kamm et al. 2020) is that model likelihoods are calculated from element-wise products of measures of deviations between data and model SFSs, thus making the likelihoods dependent on the number of elements of the SFS. This strategy inhibits comparison of models with different numbers of contemporary populations, whose corresponding SFSs have different numbers of dimensions (i.e., one dimension per population) and thus different numbers of elements. We overcome this limitation by introducing collapsed log-likelihood (CLL), in which direct comparison is enabled by “collapsing” the additional populations of higher-dimensional SFSs such that all SFSs to be compared have identical minimal shapes. For example, in order to compare three-population models of Europe that include the putative overlap zone to two-population models of Europe, we independently fit models, then “collapse” the data and model SFSs of the three-population models by summing over the axis representing the overlap zone in order to yield a 2D-SFS with the same shape as the SFSs in the two-population models, and then re-calculate the log-likelihood of the collapsed data given the collapsed model SFS in order to achieve the CLL. This method was replicated by collapsing the “Southeast” population in order to compare two- and one-population models of the “mainland” region and then by collapsing the “Latin America” population in order to compare two- and one-population models of the “Americas” region. Simulated validation of CLL as a powerful statistic for selection between models of different dimensions can be found at **Text S3**.

Replicable fitting of each model necessitated thousands of replicate runs of *moments* inference through several rounds of manual adjustment of parameter space boundaries, optimization algorithms, and other optimization parameters. The general workflow for each model fit involved initially searching enormous parameter spaces (i.e., spanning orders of magnitude in each parameter’s dimension) with the Nelder–Mead algorithm (Nelder and Mead 1965), then performing targeted searches with the BFGS algorithm (Fletcher 1987) until several runs were found to have non-randomly converged to the same point in parameter space.

To validate model likelihoods and parameter estimates, we employed a jackknifing strategy, in which, for 40 replicates for each model fit to each region, we randomly removed one sample from each population. We then calculated 95% confidence intervals as being between the second-least and second-greatest values for each estimate among each set of 40 replicates. The hypothesis tests that we reported as being performed “on model likelihoods” in the Results section are comparisons of sets of 40 CLLs of model fits to jackknife replicates.

### Linear admixture modeling and *f*_3_ analysis

We estimated the proportion of African and European admixture in North and South America, as well as Australian samples using a linear regression framework (Alkorta-Aranburu et al. 2012; Bergland et al. 2016). We modeled allele frequencies in each “admixed population” (i.e., North America, South America, Australia) as a linear combination of the two “ancestral populations” (i.e., Europe and Africa) using an intercept-free linear model:

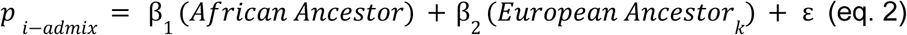

where *p_i-admix_* is a vector of allele frequencies composed of 5,000 randomly sampled SNPs across autosomes in the *i^th^*admixed sample, β_1_ represents the proportion of African ancestry and β_2_ represents the proportion of European ancestry. The model is iterated over every *k*^th^ sample from Europe and we used a sample from Zambia (sample Id = ZM_Sou_Sia_1_2010-07-16) to represent the African ancestor. We report the mean ancestry coefficients for each admix sample as the mean of β_1_ for all iterations of European ancestors. For these admixture analyses we omitted the “collapsed samples” from the (Fournier-Level et al. 2019) dataset. We performed this analysis on the entire genome, as well as inside chromosomal inversions, outside of inversions, and on non-coding mutations. In total we ran 1,313,070 comparisons (all available in **Dataset S2**).

We also assessed evidence of admixture using the *f*_3_ statistic in the R package *poolfstat* (v2.3.0, Gautier et al., 2022). A significantly negative *f*_3_ for a triplet configuration of the form *f*_3_ (A;B,C) provides evidence for the target population A to originate from an admixture event between two source populations related to sampled populations B and C. We tested samples in the Americas and Australia to identify the most likely ancestral populations from Africa and Europe. For this analysis, we included 15 African populations (derived from seven countries: Cameroon, Egypt, Ethiopia, Morocco, Rwanda, South Africa, and Zambia) and all European samples as source population proxies. We used all populations in Australia and the Americas as targets of admixture.

### Population differentiation

We analyzed patterns of population differentiation across samples and clusters using the R package *poolfstat* (v2.3.0, Gautier et al., *in prep.*). This analysis was performed for 528 samples that passed quality filtering and for 9 clusters (clusters defined based on the spatial clustering using k = 4 and continent), thus excluding the *D. simulans* sample and “CN_Bei_Bei_1_1992-09-16”, on three set of polymorphisms: i) all chromosomes including heterochromatin; ii) autosomes, excluding heterochromatin; and iii) excluding heterochromatin and SNPs with MAF < 0.05. To examine pairwise population differentiation, the samples were grouped based on their spatial clusterings at *k* = 4 and *k* = 8 (*k* = 8 clustering results shown in the supplement, **Fig. S13**). The *computeFST()* function was first used to estimate the global *F*_ST_ across all worldwide samples and also within each geographical cluster using the ANOVA method (Hivert et al. 2018).

To further quantify the impact of the structuring of the genetic diversity across continents, we used a hierarchical modeling of differentiation consisting of decomposing overall *F*_ST_ (here denoted as *hF*_ST_) into an across-group (*F*_GT_) and within group (*F*_SG_) contribution (Nei 1973), as follows:

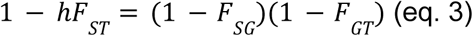

with groups of population being defined a priori (e.g., according to their continent of origin and the clustering results as we did in the present study). We estimated these statistics using the unbiased estimator developed for Pool-Seq data implemented in the *computeFST*() function of *poolfstat* (v2.3.0, Gautier et al., in prep). In addition to whole genome-estimates, window-wise hierarchical *F*_ST_ parameters were estimated across windows of 10 kb, 50 kb and 100 kb and are available in the DEST 2.0 browser.

### GIM predictive models

GIMs analyses were conducted in the R package *adegenet* v2.1.5 using discriminant analysis of the principal component (DAPC) framework (Jombart et al. 2010). While the original GIM set from DEST 1.0 consisted of 30,000 loci, here we use only 28,253 loci. This was done because some of the original markers were filtered out in the current DEST 2.0 panel. We used these markers to train the DAPC model using the sample’s state/province as the grouping prior. We retained 30 PCs from the DEST 1.0 model for the state/province model. We retained PCs based on a leave-one-out analysis that minimized the sum of squared errors (SSE) of the model. In addition, we also trained a second DEST-GIM 1.0 model using city labels (20 PCs were retained for this model; based on minimum SSE). We used 232 samples from DEST 1.0 to train the model and then predicted the provenance of all 455 new samples from DEST 2.0.

DAPC models were trained using a cross-validation routine where the data is subdivided into a training (90%) and a testing set (10%) across 30 replicates. For simplicity, we only explored the first 300 PCs across iterations. Parameters were optimized using the lowest mean square error (MSE) statistic using the *xvalDapc* function in *adegenet*. Predictive GIM models were assessed by estimating the haversine distance (*d_hav_*) between the predicted and expected latitude and longitude points. Haversine distances represent the lowest distance between two points across a spherical earth with radius of 6378.137 Km using the R package geosphere (v.1.5-14; Hijmans et al. 2022).

### Temporal genetic structure and latitudinal analysis

We assessed levels of temporal structure across DEST by estimating *F*_ST_ between samples at the same locality collected a year apart from each other. These estimates of *F*_ST_ reflect differentiation resulting from the overwintering population “bust” across one winter. We call this summary statistic “year-to-year *F*_ST_” as it captures levels of genetic variation for the population before and after a winter season. We correlated this data to latitude and performed a broken-stick regression analysis using the *segmented* (v.2.0-4) R package (Muggeo 2003).

### Scans for adaptive differentiation

We tested for adaptive differentiation at ∼908,543 SNPs that were polymorphic in a set of seasonally collected samples from across Europe (**Table S12**). First, we implemented the *BayPass* 2.4 model for adaptive differentiation using the *XtX*\* test statistic (Olazcuaga et al., 2020) while controlling for population structure using a matrix of genetic relatedness (i.e., Ω matrix). We estimated the *XtX*\* for every autosomal SNP in the genome using five independent runs of *BayPass* 2.4, and took the median value per SNP. We also generated a null distribution of *XtX*\* using the POD method outlined in Gautier (2015) and Olazcuaga et al. (2022). We generated a null distribution of *XtX*\* statistics by simulating allele frequencies for ∼9M SNPs, ten times the number of observed SNPs used in this analysis. We then generated empirical *P*-values for the observed *XtX*\* statistics by calculating the upper-tail probability of the observed data relative to the simulated POD data. We used the weighted Z analysis (wZa; Booker et al. 2024) to identify windows of signal enrichment across the genome. The wZa statistic combines the empirical *P*-values within a window for each test using Stouffer’s method (Stouffer et al. 1949) weighted by average heterozygosity. We applied this approach in a sliding window approach with a window size of 100 kb and a step size of 50 kb.

Second, we ran the *BayPass* model including both the Ω matrix as a demographic prior as well as “spring” and “fall” labels as a proxy for seasonal selection pressures. We designated the “spring” sample as the first sample within a year, and the “fall” sample as the last sample within the year. Several samples from DEST 1.0 were characterized by the collectors as “spring” or “fall”. For those samples, this label was used in the analysis. For more recent samples, including most sampled in DEST 2.0, samples are labeled as a function of date of collection. For such samples, we assigned seasonal labels by selecting the first and last sample collected in a locality within a year. For each SNP, we estimated the contrast statistics (*C*_2_) with five independent runs of *BayPass* and took the median value. To generate a null distribution of *C*_2_ statistics, we used the simulated SNP data described above, and ran *BayPass* five times. We took the median *C*_2_ of the simulated data as our null distribution, and calculated empirical *P*-values as described above. We performed a sliding window analysis of these empirical *P*-values using the wZa method.

Third, we implemented a generalized linear mixed model (GLMM) approach that is similar to that applied previously by Machado et al. (2021). We modeled allele frequency at each SNP *i* using two models :

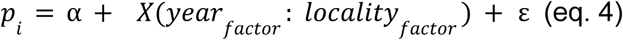

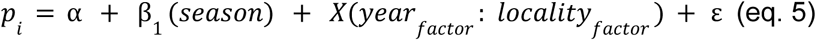

Where *p*_i_ is the allele frequency at the *i^th^*locus, α is the intercept term and β_1_ is the term associated with season, and *X* is the random effect term coded as an interaction term between the year of collection and the locality where flies were collected, ε is the binomially distributed error. We assessed the statistical significance of the seasonal β_1_ term using a likelihood ratio test between equations 4 and 5. We performed a permutation analysis following the methods outlined in (Machado et al. 2021) by shuffling the seasonal labels 100 times and rerunning the GLMM analysis for each permutation. We conducted a sliding window analysis of the GLMM.

### GO term enrichment analysis

We performed gene ontology enrichment analysis using GOWINDA v.1.12 (Kofler and Schlötterer 2012) in gene mode (with parameters: --min-genes 5 --min-significance 1 --simulations 100000) on genes located in 10 kb windows of high differentiation (*F*_GT_ > 0.2; **Table S7**), -log_10_(wZa *p*-values) > 188.96 for the *XtX*\* statistic (**Table S8)**, and -log_10_(wZa *p*-values) > 3.65 for the *C*_2_ statistic (**Table S9**), representing the 99.9th percentile from the simulated POD data (see above).

## Ethics statements

Fruit flies were collected either on public lands, where no permits are needed, or in private lands with explicit permission from the relevant stakeholders. To comply with the Nagoya protocol, material transfer agreements (MTAs) were secured here among researchers to transport fly samples (for all new samples reported here) across borders. Permit MAE-DNB-CM-2015-0030, from the Environmental Ministry of Ecuador, was obtained by Vela to collect, export and perform molecular analysis on samples.

## Author Contributions

All author contributions to this work are denoted in **Table S13**.

## Acknowledgements

We are indebted to all members of the DrosEU and DrosRTEC consortia for their support, collaboration, and for discussion over the years. DrosEU was funded by a Special Topic Networks (STN) grant from the European Society for Evolutionary Biology (ESEB). Nunez acknowledges the Henderson-Harris fellowship program at the University of Vermont, also the Vermont Advanced Computing Center (VACC; URL: https://www.uvm.edu/vacc) for providing computational resources that contributed to this publication. Bergland acknowledges Research Computing at The University of Virginia (URL: https://rc.virginia.edu) for providing computational resources and technical support that have contributed to the results reported within this publication. Coronado-Zamora and González acknowledge the Galician Supercomputing Center (CESGA), which provided access to its supercomputing infrastructure, the supercomputer FinisTerrae III and its permanent data storage system, funded by the Spanish Ministry of Science and Innovation, the Galician Government, and the European Regional Development Fund (ERDF). Gautier acknowledges the genotoul bioinformatics platform Toulouse Occitanie (Bioinfo Genotoul, https://doi.org/10.15454/1.5572369328961167E12) for providing computing resources. Obbard acknowledges Sue and Keith Obbard and Sandy Bayne for permission to collect flies on their land. Ansari acknowledges the Department of Evolution and Ecology at the University of Freiburg (Germany) for providing the necessary resources and support for sample preparations and DNA extractions. Serga acknowledges support from the PAUSE-ANR Ukraine Program. We also wish to thank Pavlo A. Kovalenko and Nadiia M. Pirko for their assistance with collecting flies in 2017-2021. Note: After 24 February 2022, no collaborative actions or exchanges have taken place within our project between Ukrainian and Russian scientists nor their institutions.

## Data availability and the new DEST 2.0 web browser

The DEST 2.0 browser is built on the latest version of JBrowse 2 (Diesh et al. 2023), an enhanced successor to JBrowse 1, which powered the original DEST 1.0 browser (Kapun et al. 2021). JBrowse 2.0 offers improved performance through a modern software architecture that supports parallel rendering of tracks and allows for the visualization of new data types, such as VCF files. Similar to the first DEST browser, it features a user-friendly data selector that facilitates the selection of the multiple population genetic metrics and statistics compiled for the DEST 2.0 release (**Fig. S16**). Additionally, the browser provides a portal for downloading allelic information and precomputed population genetics statistics in multiple formats, along with a usage tutorial featuring worked examples. Bulk downloads of all compiled tracks are available in BigWig format (Kent et al. 2010), and Pool-Seq files (in VCF format) can be accessed through a dedicated data directory. All data, tools, and supporting resources for the DEST 2.0 release, including reference tracks from FlyBase (v.6.12; Dos Santos et al. 2015), are freely available at our website (https://dest.bio). The browser operates on an Apache server running CentOS 7.2 Linux x64, powered by 16 Intel Xeon 2.4 GHz processors and 32 GB of RAM. All sequences are available on the SRA (https://www.ncbi.nlm.nih.gov/sra) at PRJNA993612. Code is available in GitHub at: https://github.com/DEST-bio/DESTv2_data_paper. All outputs from the DEST 2.0 pipeline can be found at https://dest.bio. Supplementary datasets can be found in Zenodo at https://doi.org/10.5281/zenodo.13731977.

