## Supplementary material for "Footprints of worldwide adaptation in structured populations of *D. melanogaster* through the expanded DEST 2.0 genomic resource": Fig. S

Contents:

---


**Supplementary Figures:**

Fig S1: DEST 2.0. description and quality control statistics
Fig S2: $p_n/p_s$ analysis
Fig S3: Contamination analysis
Fig S4: Variable contribution to quality control filtering
Fig S5: Recombination landscape in 75 *Drosophila melanogaster* populations
Fig S6: Variation in the recombination rate landscapes
Fig S7: Random allele PCA
Fig S8: Cosmopolitan inversions in DEST 2.0
Fig S9: Conceptual models for the *moments* analysis
Fig S10: Likelihood distributions from the *moments* analysis
Fig S11: African ancestry in the linear admixture analyses
Fig S12: $F_{ST}$ estimates with the block−jackknife resampling approach
Fig S13: $F_{ST}$ estimates over all DEST 2.0 samples
Fig S14: Signatures of local adaptation in *Cyp* genes
Fig S15: Enrichment analysis from $C_2$ SNPs
Fig S16: Browser snapshot

---

**Supplementary Tables:**

Table S1: DEST 2.0 metadata.
Table S2: Correlation between recombination rate and diversity statistics.
Table S3: Recombination and inversions analysis.
Table S4: Inversion clines analysis.
Table S5: Optimal demographic model parameters.
Table S6: Admixture (*f*3) summary analysis.
Tables S7-S9: GOWINDA results.
Table S10: Genes in the enriched GO term GO:0004022.
Table S11: Correlation between recombination rate estimations.
Table S12. Samples used for seasonal analysis of European samples.

---

**Supplementary Data:**

Dataset S1: *f*3 comparisons (1,478,000 three-population comparisons)
Dataset S2: linear admixture estimates (1,313,070 comparisons)
Dataset S3: data for seasonal analyses

---

**Supplementary Text:**

Text S1: Note effective size.
Text S2: Filtering recommendations to users.
Text S3: CLL validation.

### Supplementary Figures

**Fig S1 (figure in next page). DEST 2.0 data description.** A) Sampling effort of DEST 1.0 and 2.0. B) Number of samples collected per year across selected localities of DEST with multiyear sampling. C) Various metrics of quality control in DEST. 2.0. The names at the top of columns represent various constituent datasets: "*cville*" is the data from Virginia of Nunez *et al.* (2024), "*dest_plus*" is the added pooled samples (see main text), "*drosEU*" represents samples from the first release (DEST 1.0), "*drosEU_3*" and "*drosEU_SA*" are sampled generated for this release of the dataset, the "SA" label represents the subset of samples from South America, "*drosRTEC*" represent samples from Bergland *et al.* (2016) and "*dgn*" are samples from the Drosophila Genome Nexus (Lack *et al.* 2016).

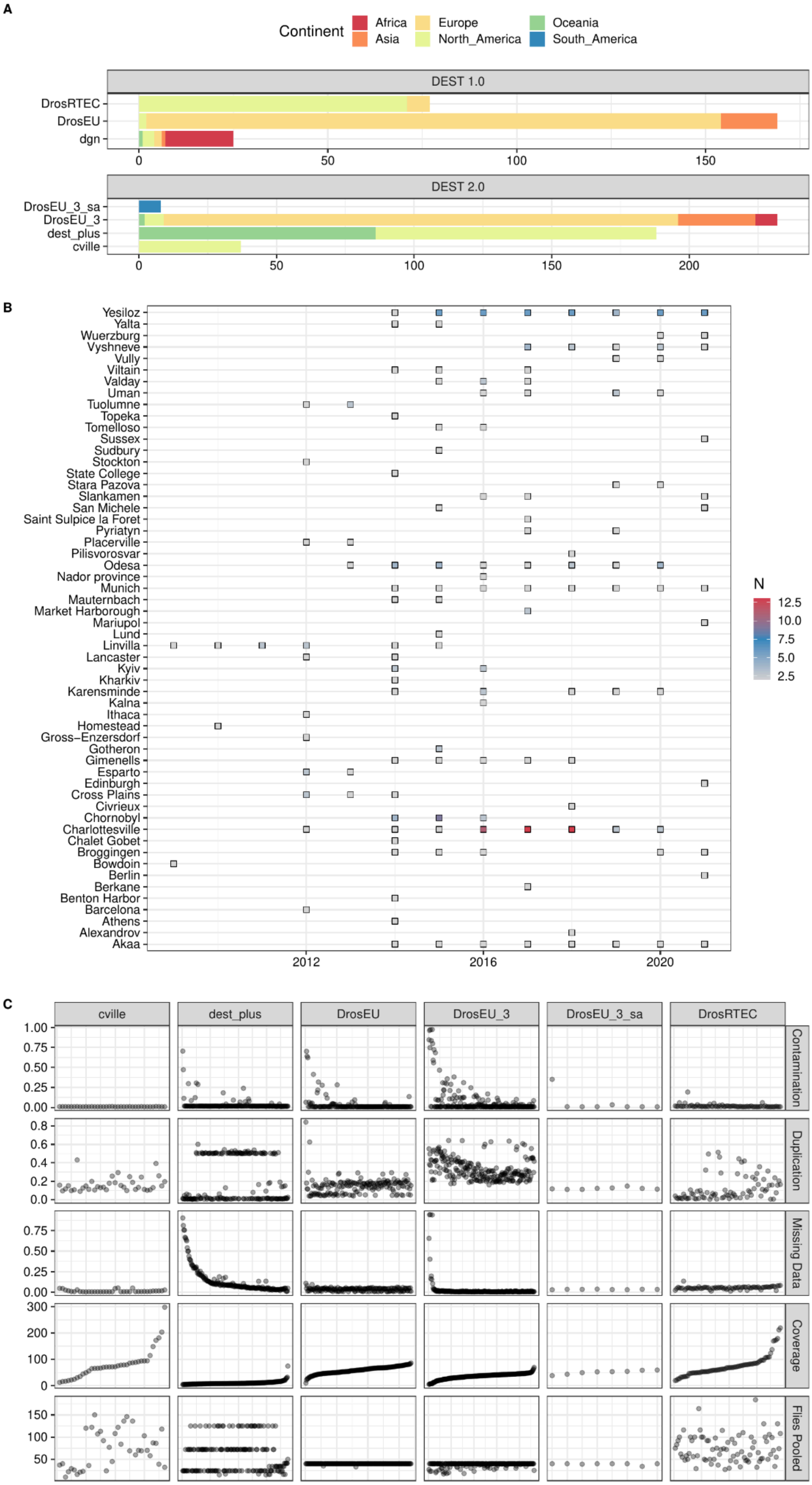
A
Continent
Africa
Asia
Europe
North_America
Oceania
South_America
DEST 1.0
DrosRTEC
DrosEU
dgn
0
50
100
150
DEST 2.0
DrosEU_3_sa
DrosEU_3
dest_plus
cville
0
50
100
150
200
B
Yesiloz
Yalta
Wuerzburg
Vyshneve
Vully
Viltain
Valday
Uman
Tuolumne
Topeka
Tomelloso
Sussex
Sudbury
Stockton
State College
Stara Pazova
Slankamen
San Michele
Saint Sulpice la Foret
Pyriatyn
Placerville
Pilisvorosvar
Odesa
Nador province
Munich
Mauternbach
Market Harborough
Mariupol
Lund
Linvilla
Lancaster
Kyiv
Kharkiv
Karensminde
Kalna
Ithaca
Homestead
Gross−Enzersdorf
Gotheron
Gimenells
Esparto
Edinburgh
Cross Plains
Civrieux
Chornobyl
Charlottesville
Chalet Gobet
Broggingen
Bowdoin
Berlin
Berkane
Benton Harbor
Barcelona
Athens
Alexandrov
Akaa
2012
2016
2020
N
12.5
10.0
7.5
5.0
2.5
C
cville
dest_plus
DrosEU
DrosEU_3
DrosEU_3_sa
DrosRTEC
Contamination
Duplication
Missing Data
Coverage
Flies Pooled
1.00
0.75
0.50
0.25
0.00
0.8
0.6
0.4
0.2
0.0
0.75
0.50
0.25
0.00
300
200
100
0
150
100
50

**Fig S2.** Genome-wide $p_N/p_S$ ratios and the $\log_{10}$-scaled number of private SNPs for all Pool-Seq samples. Similarly to our analysis in the first release of DEST (i.e., Kapun *et al.* 2021), we highlight outlier samples in red that are characterized by unusually high values for both numbers of private SNPs and $p_N/p_S$. The dashed black lines indicate both statistics' 95% confidence limits (average + 1.96 SD). The vertical green dashed line highlights the empirical estimate of $p_N/p_S$ calculated from individual sequencing data of the DGRP freeze2 data set. The green diamond represents the corresponding value of the DGRP population, which was pool-sequenced as part of the DrosRTEC data set (US_Nor_Ral_2_2003-07-01 in DEST 2.0 nomenclature).

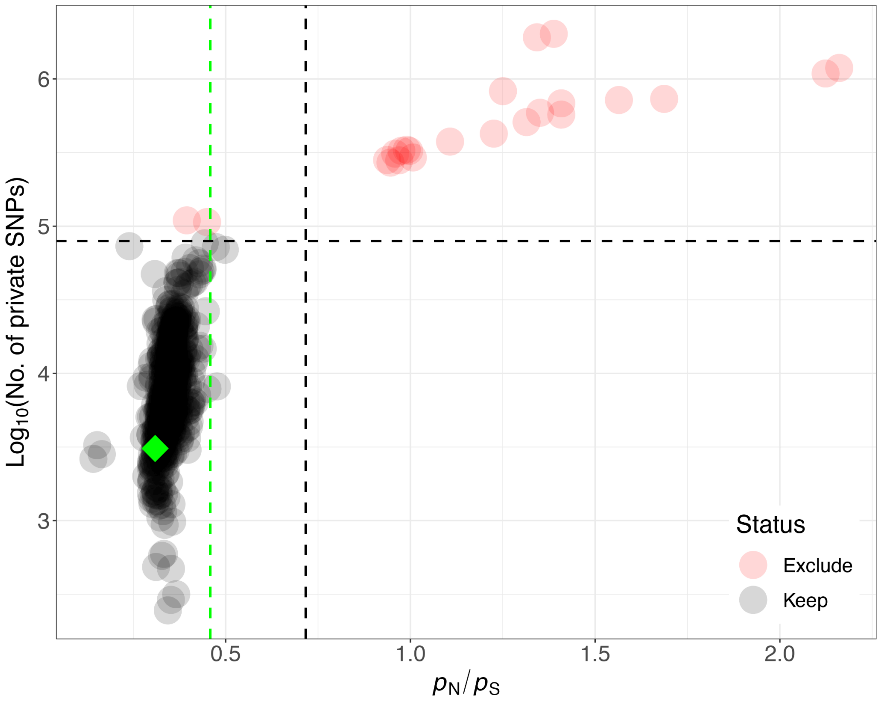

**Fig S3.** Contamination analysis. A) shows the relationship between the estimated amount of contamination of a simulated dataset vs. the "real" level of contamination from the in-silico mixing of inbred *D. simulans* or *D. melanogaster* genomes. The two colors represent the approaches to detect contamination (see Main Text). B) Correlation of *D. simulans* contamination estimates across both approaches (i.e., *k*-mer vs. competitive mapping) in the DEST data. C) Estimated levels of *D. simulans* contamination across sample sets in DEST 2.0. D) Estimated contamination levels derived from any taxa with >1% representation based on the *k*-mer approach. The arrows in panels B and D point to the sample with a low amount of contamination by *D. serrata*. That sample has higher estimates of *D. simulans* contamination using the competitive mapping approach.

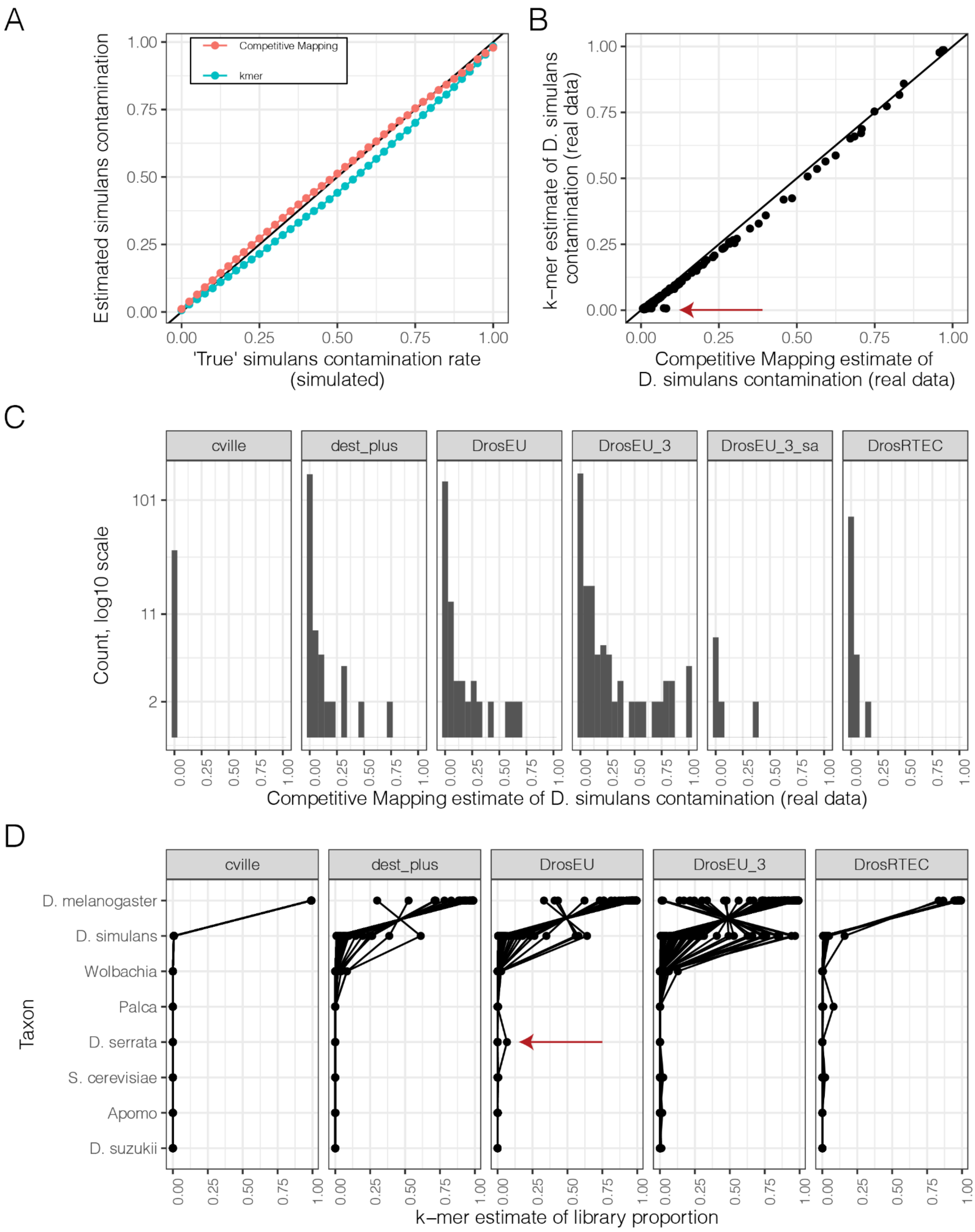

**Fig S4.** DEST quality control. Variables (i.e., QC metrics) contribution to the PCA shown in panel 2A of the Main Text. The color indicates the contribution to sample projections.

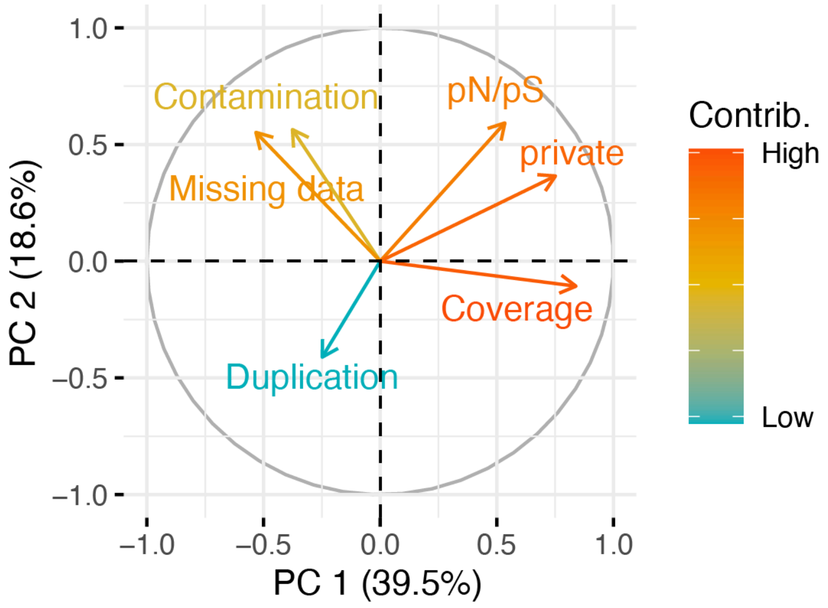

**Fig S5.** Recombination landscape in 75 *Drosophila melanogaster* populations (gray lines). Average (black line) and 95 % confidence interval (light blue ribbon; calculated using a normal distribution assumption) are shown for each chromosome. Recombination rate was estimated for 100 kb non-overlapping windows in autosomes, and 200 kb windows in the X chromosome. The boundaries of the main inversions are identified by red dotted lines.

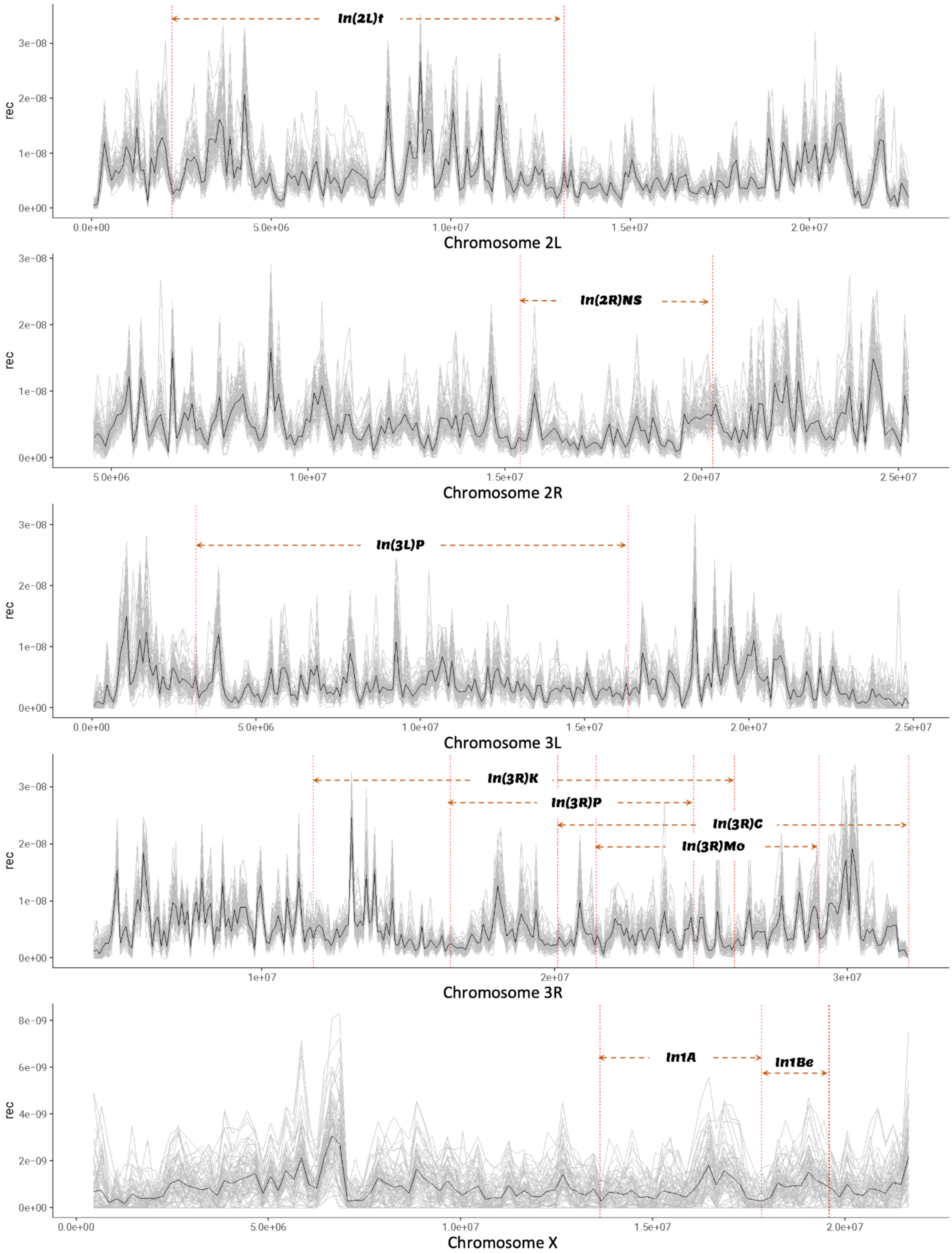

**Fig S6.** Variation in the recombination rate across 75 *Drosophila melanogaster* populations. The PCA analysis is based on the genome-wide recombination rates estimated in 200 kb non-overlapping windows along the genome. PCA in panels A and B are based on raw recombination rate values, while PCA in panels C and D are based on relative recombination values (raw value/chromosome average). Group coding is based on clustering at $k = 4$, and sample codes are as in Table S2.

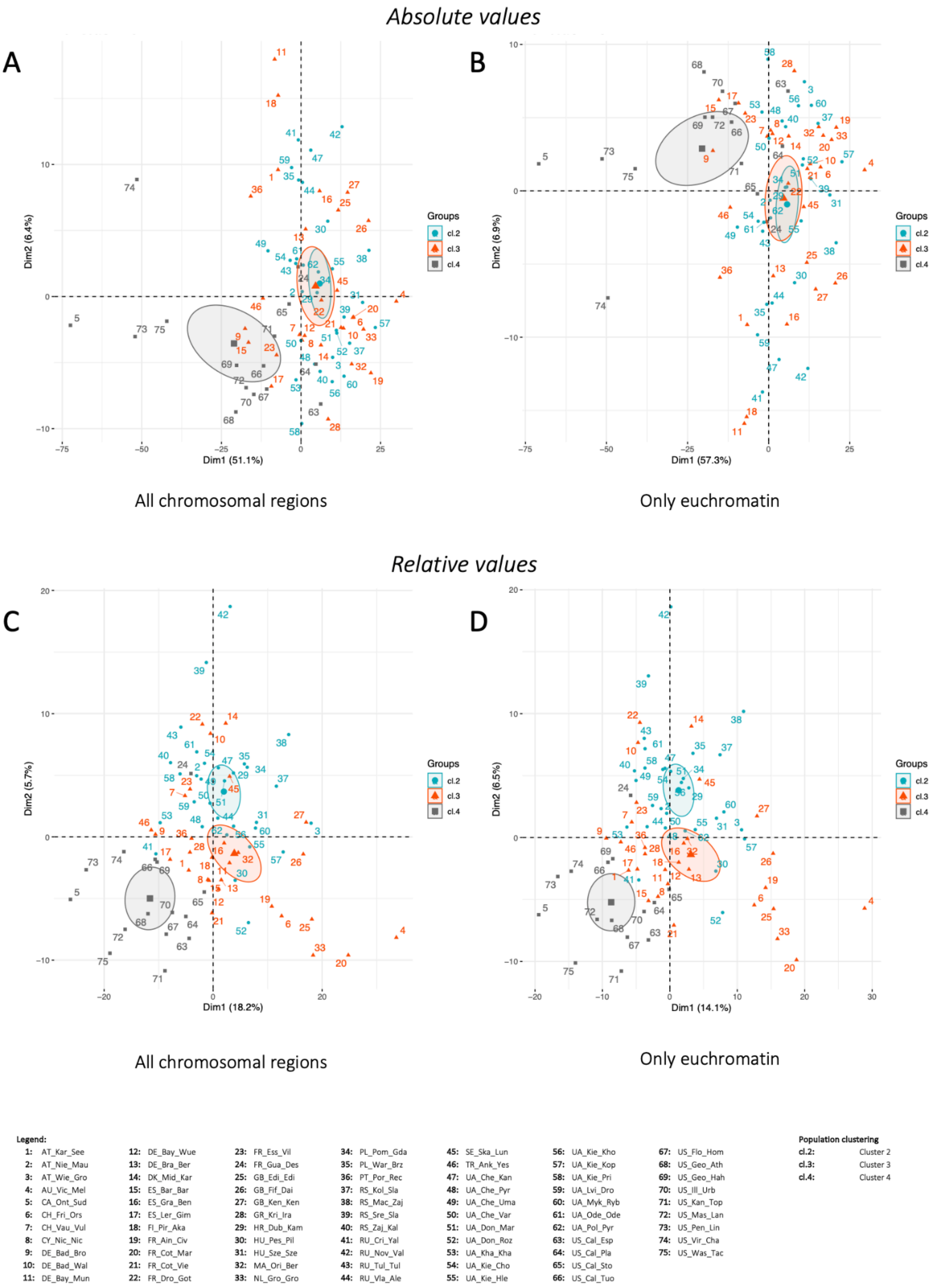


**Fig S7. Principal component analysis and projections using a random allele approach.** A) PCA projections showing PCs 1 and 2. Analyses were done for each autosomal chromosome arm, and all arms were combined. B) PCA projections showing PCs 1 and 3. Percentage of variance explained for PC1, PC2 and PC3, respectively: 2L: 1.97%, 0.98% and 0.49%; 2R: 2.06%, 1.06% and 0.48%; 3L: 2.23%, 1%, 0.48%; 3R: 2.48%, 1.12%, 0.57%; X: 4.02%, 2.3%, 0.77%. To compare both

PCAs, we calculated the Euclidean distance between all points in the PC1 and PC2, and PC1 and PC3 of the PCAs using the dist() function from the stats package and performing a canonical correlation with the cancor() function of the stats package. The correlation was 0.99 for PC1 and PC2 and 0.91 for PC1 and PC3. Therefore, variations in pool coverages and sizes impact the pattern observed for the PCA. C) Correlation coefficients (confidence intervals) among principal components and latitude and longitude, across chromosomes.

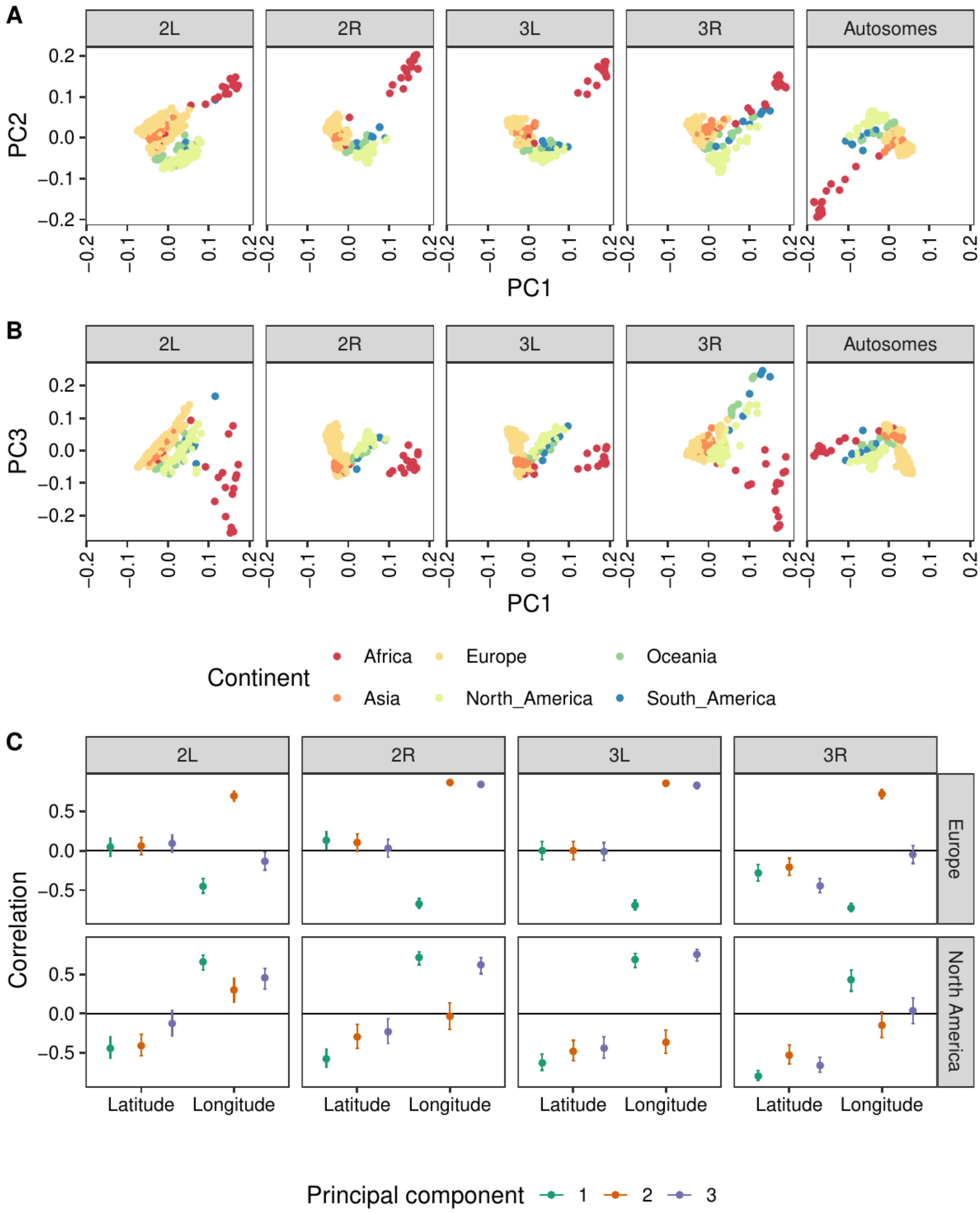

**Fig S8.** Frequencies (see color legend) of the four most common cosmopolitan inversion polymorphisms in worldwide populations estimated from inversion-specific markers SNPs. Estimates from the present data are indicated by squares and previous estimates from Kapun & Flatt (2019) are shown as circles.

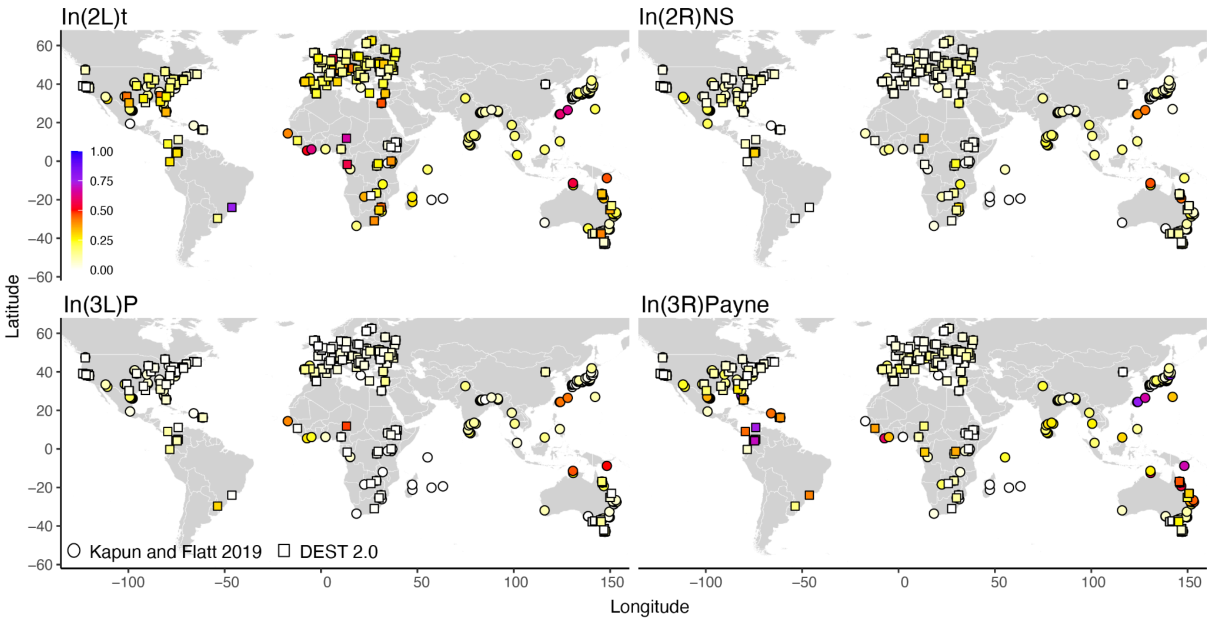

**Fig S9.** Sketches of the *demes*-type demographic models fit subsets of the variant data with *moments*. Each circle represents a population, and a circle featuring an "X" over it represents an extinct ancestral population. Solid lines represent direct descendants, and dashed lines represent descendants via admixture. Migration is not explicitly shown.

| Explanation of demographic models fit in this analysis | | | |
|---|---|---|---|
| Model | # of pops. | # of params | Description |
| One-pop. | 1 | 2 | An ancestral population instantaneously changes at time $T$ to size $N$. |
| Split | 2 | 4 | An ancestral population splits into two derived populations ($N_1$ and $N_2$) at time $T$, followed by constant symmetric migration ($m$). |
| Two-splits | 3 | 8 | A single ancestral population splits into two derived populations ($N_1$ and $N_{int}$) at time $T_1$, followed by constant symmetric migration ($m_2$). Then, at time $T_2$, the intermediate (int) population splits into two derived populations ($N_2$ and $N_3$), followed by constant symmetric migration among all three populations ($m_3$). |

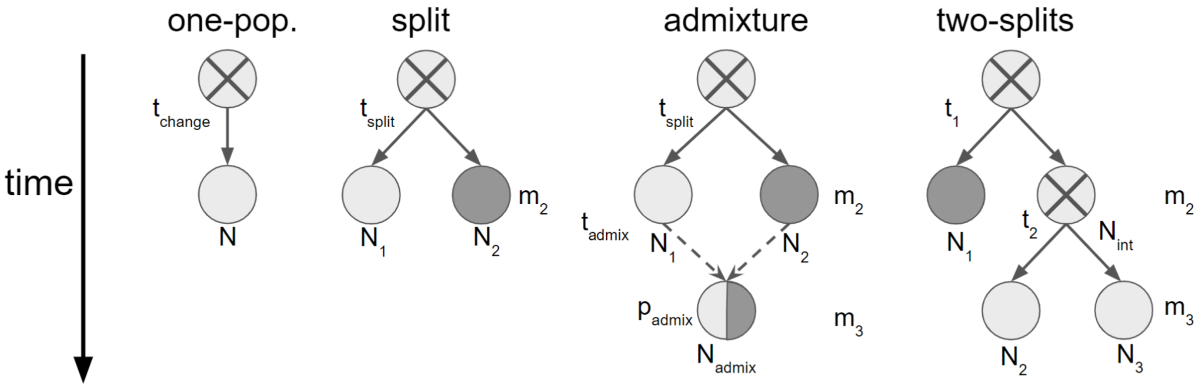

**Fig S10.** Violin plots of log-likelihoods of optimal model fit to jackknife replicates (*n* = 40) of variant data within each subset of the data (regions). Collapsed log-likelihood enables direct comparison of models that include additional populations, so, in models with the minimal number of populations for a given region (namely "one-population" for the mainland [A] and the Americas [B] and "split" for Europe; [C]), collapsed log-likelihood equals log-likelihood. "One-population" was selected as the optimal model for both mainland and the Americas by pairwise comparison with Wilcoxon signed-rank tests of the sets of (collapsed) log-likelihoods. "split" was selected as the optimal model for Europe by the Kruskal–Wallis *H* test ($P = 8.7 \times 10^{-28}$) followed by six Benjamini–Hochberg-corrected Dunn's tests of "split" against each other model. The regions are described in the Results and the models are described in the Methods sections of the main text. *P*-values not given here can be found in the Results section of the main text.

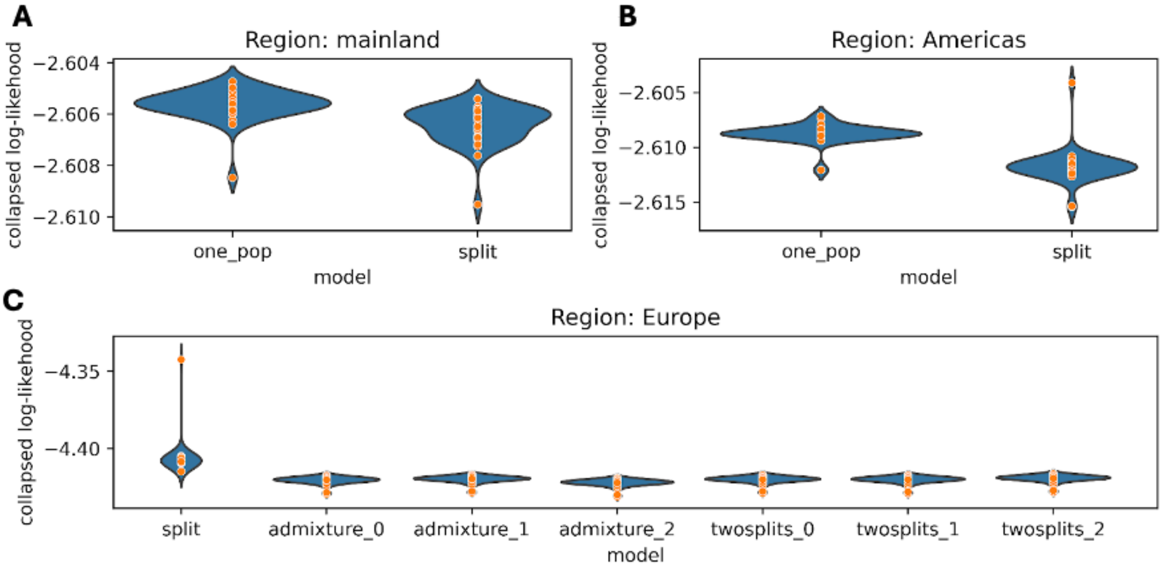

**Fig S11.** African ancestry results in the context of our clustering analyses (*k*=8). A) All North American clusters of k=8 can be differentiated as a function of their African ancestry. The Caribbean cluster (cluster $5_{k=8}$, green dots in Fig. 4D) is defined by high levels of African admixture (mean $\beta_{African}$ = 0.36). Cluster $6_{k=8}$ comprises pools from the southern seaboard of eastern North America (i.e., Virginia, Georgia, South Carolina) and has a mean $\beta_{African}$ = 0.23. All other samples from continental North America (cluster $4_{k=8}$) have lower relative levels of African ancestry (mean $\beta_{African}$ = 0.19). These results confirm that levels of African admixture are a key driver of the population structure of *D. melanogaster* outside of Africa and Europe. B) Proportion of significant admixture tests (see color gradient) when the population of interest is used as the African parent.

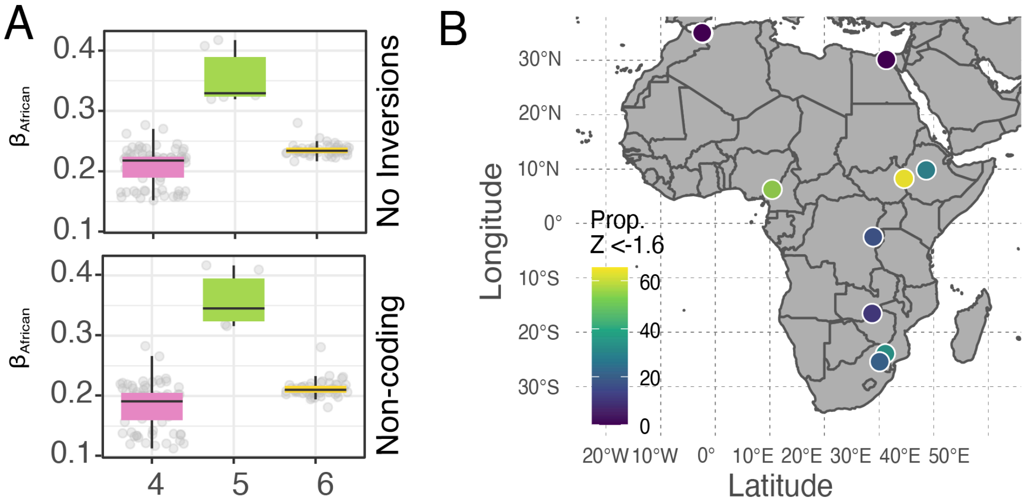

**Fig S12. $F_{ST}$ analyses. A)** Standard $F_{ST}$ calculations in DEST 2.0 using a 50,000 block−jackknife resampling approach. As Fig. 6A, but only considering euchromatin (left) or euchromatin and SNPs with MAF < 0.05 (right). The color indicates autosomes (blue) and X chromosomes (orange). **B)** Hierarchical $F_{ST}$ calculations in DEST using a 50,000 block−jackknife resampling approach. This is like Fig. 6A (in the main text), yet it only considers euchromatin (left) or euchromatin and SNPs with MAF < 0.05 (right).

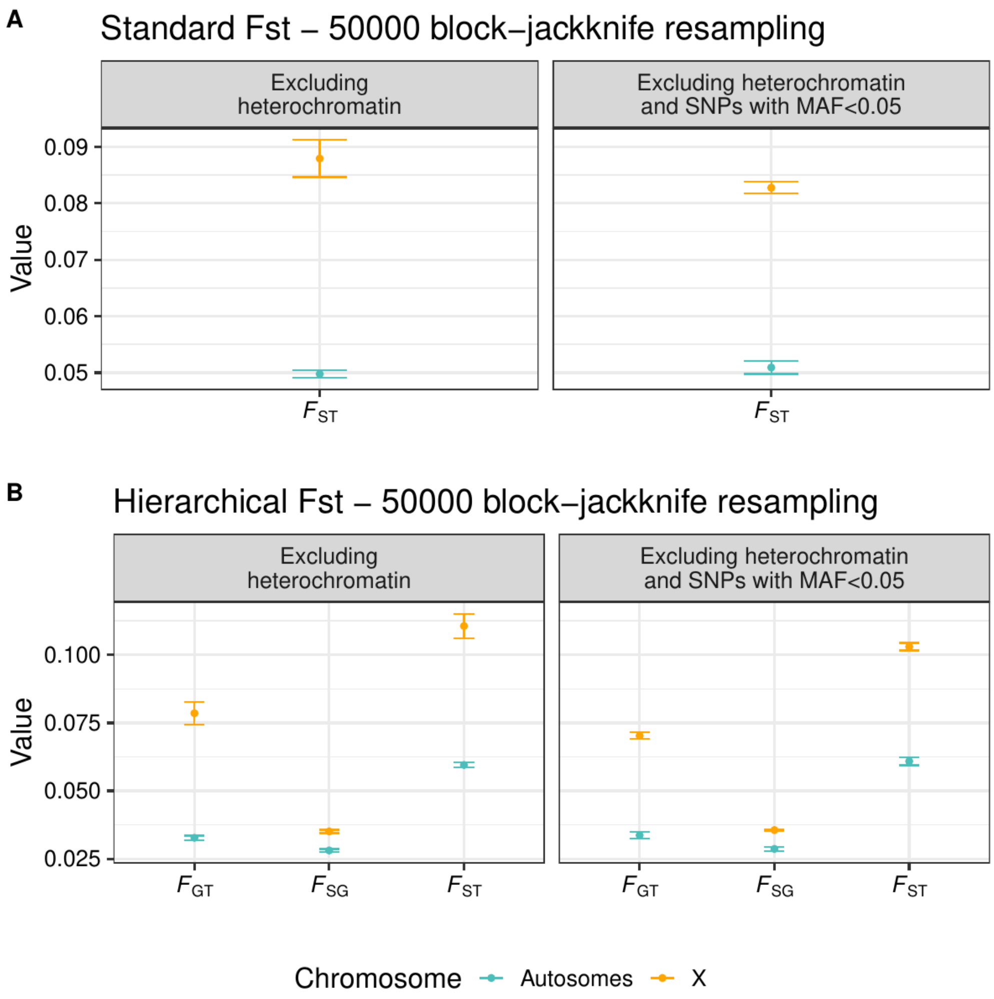

**Fig S13.** **A)** Standard $F_{ST}$ estimates over all DEST samples and their 95% CI (corresponding to +/-1.96 s.e. estimated using a block−jackknife with blocks of 50,000 consecutive SNPs) for samples across clusters from the $k = 8$ analysis (orange = X chromosome; green = autosomes), including SNPs in heterochromatin. **B)** Pairwise comparisons between cluster-continents estimated as $F_{GT}$ (under $k$ = 8) result in a heatmap. Only autosomes are considered. These figures are equivalent of the main text to **Figures 6C** and **6B**, respectively.

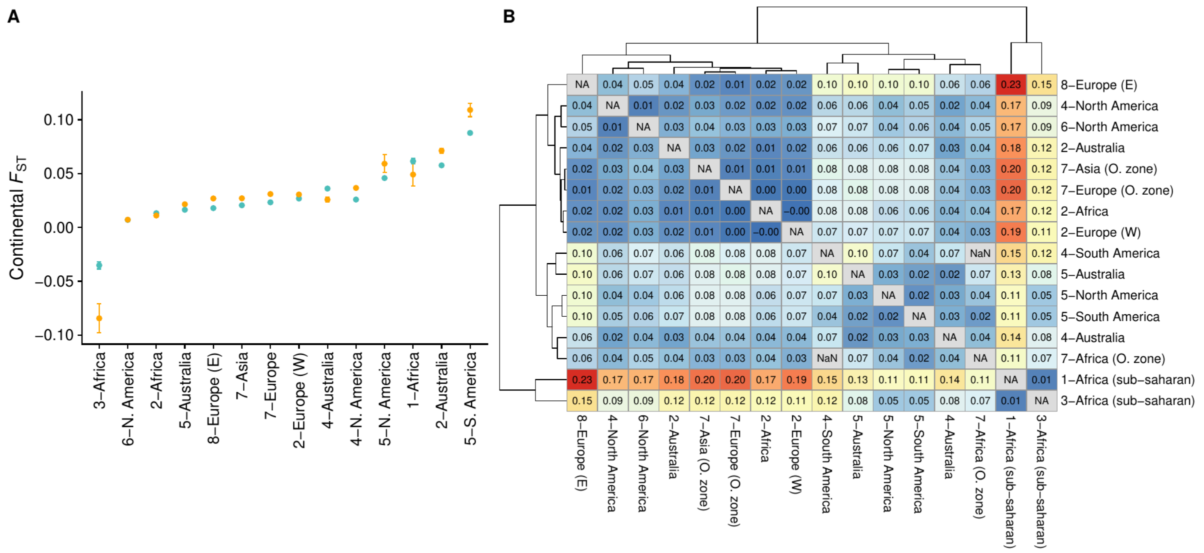

**Fig S14.** Pairwise differentiation was measured with $F_{GT}$ of chromosome 2R between North America (cluster 4) and West Europe (cluster 3), North America (cluster 4) and South America (cluster 4), and South America (cluster 4) and West Europe (cluster 3). Lastly, a Zoom-in on the region, showing a cluster of *Cyp* genes is shown.

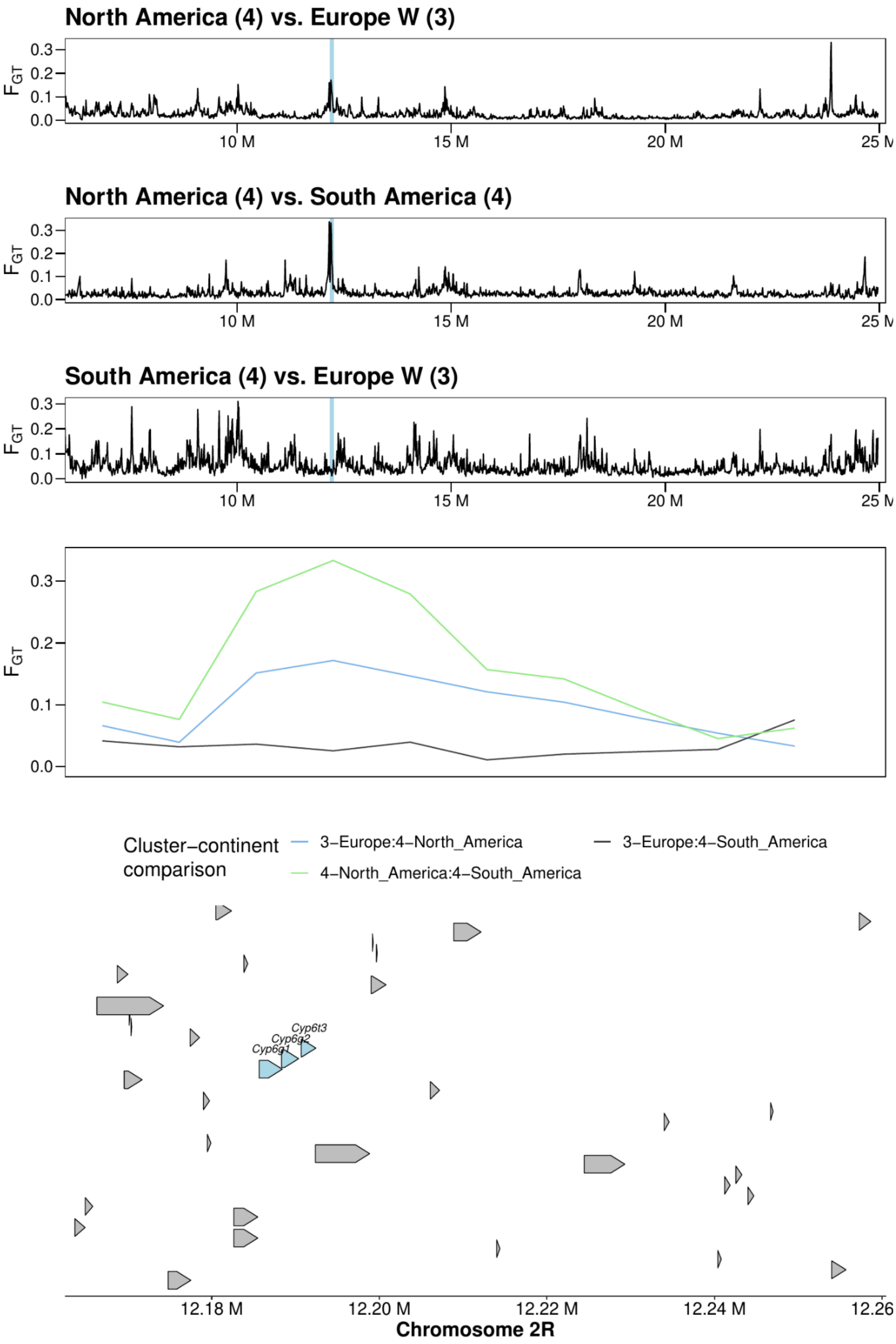

**Fig S15 Enrichment analysis from c2 SNP compared to other seasonal Spring vs. Fall analyses. (A)** Enrichment analyses of the top $1.0x10^{-4}$ *c2* SNPs among the top seasonal SNPs of Bergland et al. 2014 (FDR < 0.3 as reported in their paper). **(B)** Same as A but with the top 1% seasonal SNPs in Machado et al. 2021. The plot shows $log_2$ transformed odds ratio with 95% confidence intervals. Significant observations display a * symbol.

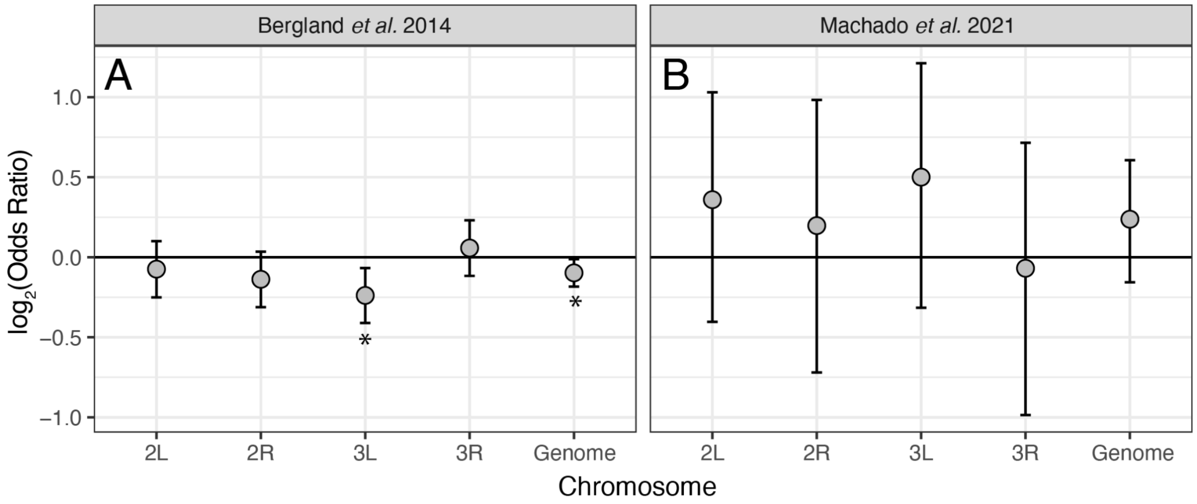

**Fig S16.** Snapshot of the new DEST 2.0 browser. It shows a region containing a cluster of P450 genes in chromosome 2R (*Cyp6g1*, *Cyp6g2* and *Cyp6t3*). Tracks activated include the Gene annotations, Inversions, and Transposable elements (corresponding to reference genome r6.12); the recombination in cM/Mb retrieved from Comeron et al. (2012), and the population differentiation between Europe (cluster 2) and North America (cluster 4), estimated with Fgt. The left menu is the new track selector of JBrowse 2, displaying all track categories available in DEST 2.0.

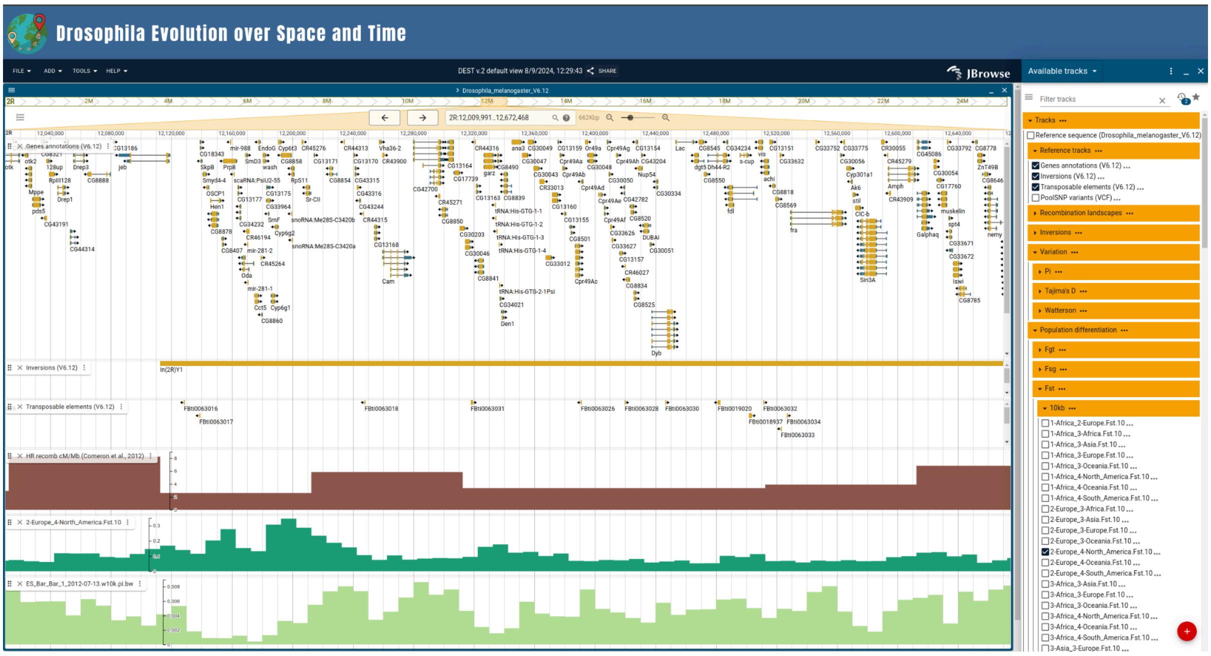

**Supplementary Data**
Supplementary Data is available in Zenodo at
https://doi.org/10.5281/zenodo.13731977

Nunez, J. C. B. (2024). Supplementary Data of DEST 2.0 [Data set]. Zenodo.
https://doi.org/10.5281/zenodo.13731977

**Supplementary Text**
Supplementary Text is available at the Journal website
