## Supplementary material for "Footprints of worldwide adaptation in structured populations of *D. melanogaster* through the expanded DEST 2.0 genomic resource": Text S1

Following GAUTIER *et al.* (2013), in a Pool-Seq experiment, the effective diploid sample size $n_e$ (or effective coverage) of the DNA pool can be defined as the number of individually genotyped (haploid) individuals that would show the same amount of variance in the allele frequency estimate (see also KOLACZKOWSKI *et al.* (2011) and FEDER *et al.* (2012)). For an (arbitrarily) chosen allele at a given SNP, let $p$ represent the population frequency and $y_e$ the (allele) count among a sample of $n_e$ individuals. The binomial sampling variance of the standard and unbiased population allele frequency estimate $\hat{p}_{ind} = \frac{y_e}{n_e}$ from allele count data is thus simply:

$$\mathrm{V}\left(\hat{p}_{ind}\right) = \frac{1}{n} p \left(1 - p\right) \tag{1}$$

Let $r$ be the number of reads (i.e., read count) carrying the allele, and $c$ be the total number of reads (coverage) observed in the sequencing data for the pool. As shown previously, $\hat{p}_{pool} = \frac{r}{c}$ provides an unbiased estimator of $p$ from Pool-Seq data under a wide range of experimental conditions (GAUTIER *et al.* 2013). Assuming equal contribution of individuals to the pool sequences (see GAUTIER *et al.* (2013) for more complex modeling), i.e. $r \sim Bin(\frac{y}{n_h}, c)$ (where $y$ is the reference allele count among individuals in the pool sample and $n_h$ is the haploid sample size of the pool), the variance of this estimator can be easily derived and expressed as a function of $n$ and $c$ since:

$$\begin{aligned}
\mathrm{V}\left(\hat{p}_{pool}\right) = \frac{1}{c^2}\mathrm{V}\left(r\right) &\equiv \frac{1}{c^2}\left(\mathrm{V}_Y\left(\mathrm{E}\left(r \mid y\right)\right) + \mathrm{E}_Y\left(\mathrm{V}\left(r \mid y\right)\right)\right) \\
&= \frac{1}{c^2}\left(\mathrm{V}_Y\left(c\frac{y}{n_h}\right) + \mathrm{E}_Y\left(c\frac{y(n_h - y)}{n^2}\right)\right) \\
&= \frac{1}{c^2}\left(\frac{c^2}{n_h^2}\mathrm{V}_Y\left(y\right) + \frac{c}{n_h}\mathrm{E}_Y\left(y\right) - \frac{c}{n_h^2}\mathrm{E}_Y\left(y^2\right)\right) \\
&= \frac{1}{n_h^2}n_h p(1-p) + \frac{1}{n_h c}n_h p - \frac{1}{n_h^2 c}n_h p(1 - p + n_h p) \\
&= \frac{1}{c n_h^2}\left(c n_h p(1-p) + n_h^2 p - n_h p(1 - p + n_h p)\right) \\
&= \frac{1}{c n_h^2} n p(c + n_h - 1)(1-p) \\
&= \frac{c + n_h - 1}{c n_h} p(1-p) \qquad (2)
\end{aligned}$$

Equating the equations 1 and 2, we get $\frac{1}{n_e} = \frac{c+n_h-1}{cn_h}$ and

$$n_e = \frac{cn_h}{c + n_h - 1} \tag{3}$$

Note that the equation 3 differs from FEDER *et al.* (2012), presumably due to a typo in the later manuscript (i.e., the term -1 in the numerator should actually be in the denominator). More generally, if the haploid sample size is much larger than the coverage (i.e., $n_h \gg c$), $n_e \simeq \frac{cn_h}{n_h} = c$. Conversely, if the coverage is much larger than the haploid sample size (i.e., $c \gg n_h$), $n_e \simeq \frac{cn_h}{c} = n_h$.

To account for the special case where multiple Pool-Seq data from the same population (but each containing different individuals) are collapsed into a single one, as in the data from FOURNIER-LEVEL *et al.* (2019), we can generalize this estimator of $n_e$ to account for variation across pools. Following the notations above, let $r_i$ be the number of reads carrying the reference allele, and $c_i$ be the total number of reads (coverage) observed in the sequencing data for the $i$th pool (out of $I$ pools), which has a haploid sample size of $n_i$ individuals and (unobserved) allele count $y_i$. The allele frequency estimator obtained from the collapsed data set can then be defined as $\hat{p}_{pool} = \frac{\sum_{i=1}^{I} r_i}{\sum_{i=1}^{I} c_i}$. This estimator is unbiased:

$$\mathrm{E}\left(\hat{p}_{pool}\right)=\frac{1}{\sum\limits_{i=1}^{I}c_i}\sum_{i=1}^{I}\mathrm{E}\left(r_i\right)=\frac{1}{\sum\limits_{i=1}^{I}c_i}\sum_{i=1}^{I}\mathrm{E}_Y\left(\mathrm{E}\left(r_i \mid y_i\right)\right)=\frac{1}{\sum\limits_{i=1}^{I}c_i}\sum_{i=1}^{I}\frac{c_i}{n_i}\mathrm{E}\left(y_i\right)=\frac{1}{\sum\limits_{i=1}^{I}c_i}\sum_{i=1}^{I}c_i p=p \tag{4}$$

and its variance is equal to:

$$\mathrm{V}\left(\hat{p}_{pool}\right)=\frac{1}{\left(\sum\limits_{i=1}^{I}c_i\right)^2}\mathrm{V}\left(r_i\right)=\frac{1}{\left(\sum\limits_{i=1}^{I}c_i\right)^2}\sum_{i=1}^{I}\left(\frac{c_i}{n_i}(c_i+n_i-1)\right) \tag{5}$$

This leads to a more general formula for $n_e$ (note that $I=1$ leads to the equation 3):

$$n_e=\frac{\left(\sum\limits_{i=1}^{I}c_i\right)^2}{\sum\limits_{i=1}^{I}\left(\frac{c_i}{n_i}(c_i+n_i-1)\right)} \tag{6}$$

These formulas can be checked using simple Monte Carlo simulations, as implemented in the *check_estimator* function provided below.

Figure 1 shows a comparison of the different estimators, including the erroneous one by Feder *et al.* (2012), with simulated values under different scenarios. For a single pool (Figure 1A and 1B), the different estimators are accurate, with Feder *et al.* (2012) estimator being only slightly biased downward. This was expected from the equations given the range of haploid samples and coverage considered. Figures 1C and 1D demonstrate the importance of properly accounting for between-pool variation in the estimation of $n_e$ when collapsing samples from multiple pools, especially when haploid sample sizes are variable. In fact, using the naive estimator based on eq. 3 with the collapsed values (i.e., with $c=\sum\limits_{i=1}^{I}c_i$ and $n_h=\sum\limits_{i=1}^{I}n_i$) results in substantial upward bias.

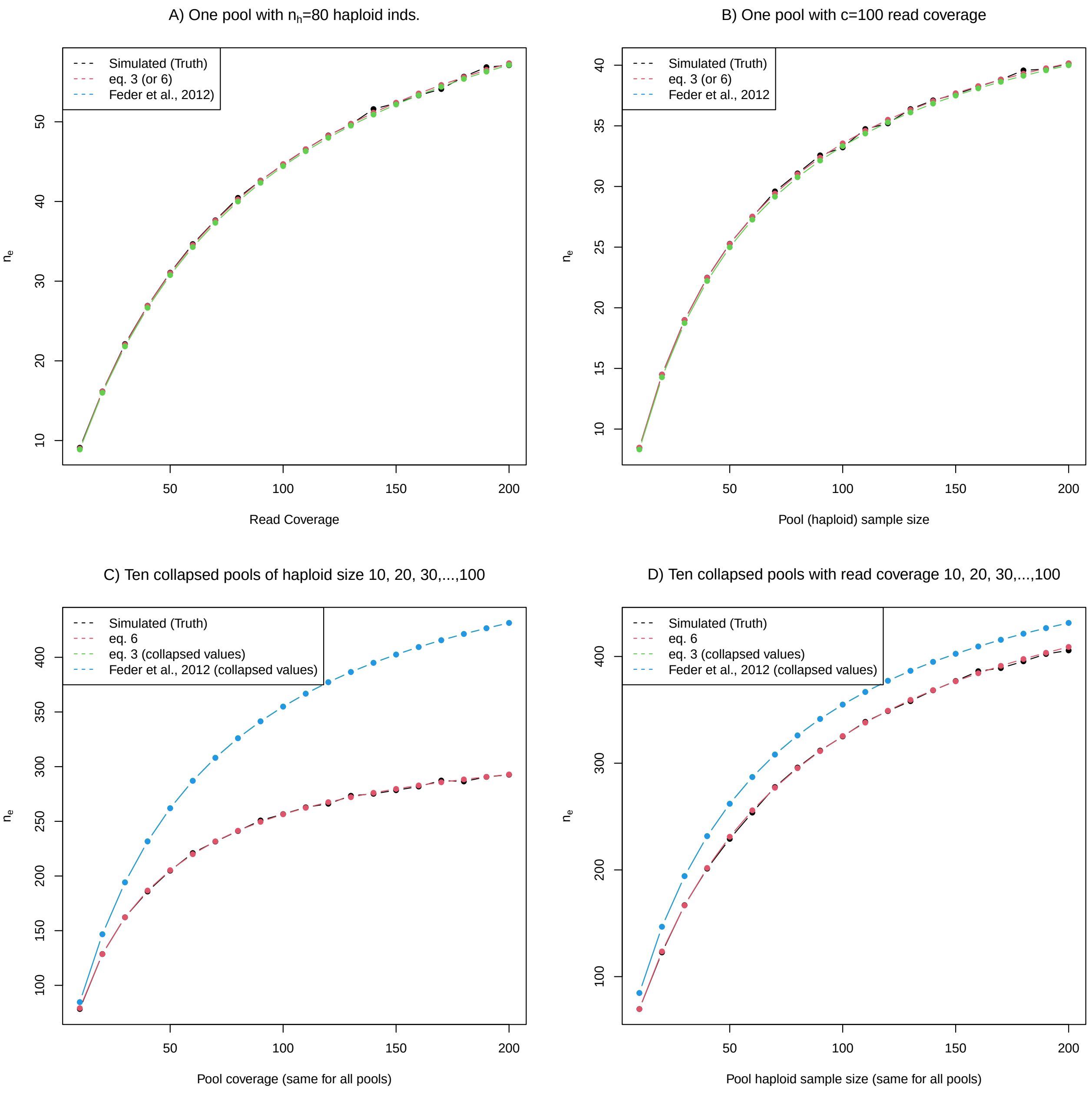


Figure 1: Comparison of simulated (n=100,000 data sets per condition) and predicted values under different scenarios involving a single pool (A and B) or ten pools from the same populations (C and D). The simulated reference allele frequency is p=0.25 in all simulations.
