## Supplementary material for "Footprints of worldwide adaptation in structured populations of *D. melanogaster* through the expanded DEST 2.0 genomic resource": Text S2

**Data filtering procedures and recommendations for users**

To ensure the highest possible quality of the data, we explored the quality of single nucleotide polymorphism (SNP) calling in our dataset by estimating the ratio of non-synonymous SNP calls ($p_N$) relative to that of synonymous SNP calls ($p_S$), i.e., the $p_N/p_S$ statistic. This statistic relies on the expectation that, on average, protein-coding regions should experience stronger purifying selection against non-synonymous changes as compared to synonymous changes, thus it can be used to assess the degree of erroneous calls across libraries (Fig. S2). Next, we estimated the levels of non-*D. melanogaster* contamination and developed a data-driven approach to filter out low-quality samples across libraries. We assessed non-*D. melanogaster* contamination in a two-pronged approach. First, we used competitive mapping between the reference genomes of *D. melanogaster* and *D. simulans* as in the first DEST release (Kapun et al. 2021) by comparing the proportion of reads that mapped to each genome. We then determined contamination using a *k*-mer approach as described by Gautier (2023). This is a computationally efficient approach to quantify levels of multispecies contamination (i.e., contamination by congeners other than *D. simulans*) without the need to map reads to a hologenome. We assessed the accuracy of each method for estimating the proportion of *D. melanogaster* and *D. simulans* using *in silico* pooled samples. Both methods estimate the contamination rate with high accuracy ($r$ > 99.95%; Fig. S3) although the competitive read mapping has lower root mean square deviation (RMSE) than the *k*-mer based (0.133 vs. 0.195, respectively). Next, we calculated contamination rates using the DEST 2.0 data. Most samples had negligible contamination rates, although there were notable exceptions, with some samples composed of nearly 100% *D. simulans* (Figs. S3B-C). Interestingly, the sample with the greatest discrepancies between the competitive read mapping-based and k-mer-based methods were those with non-zero rates of contamination with *D. serrata* (Fig. 3; Figs. S3B-D).

To make filtering recommendations, we performed PCA on several quality control descriptors across all samples. These quality control metrics include the level of *D. simulans* contamination, the rate of PCR duplication, the fraction of missing data, coverage, the number of private SNPs, and the $p_N/p_S$ ratio. Using the first two PCs of the QC analyses (39.5% and 16.6% of variance explained, respectively) we clustered samples based on their quality control properties (Figs. 1C and S4). The PCA allowed us to divide the samples into four basic recommendation categories based on *k*-means clustering: samples that passed all filters (purple triangles; Fig. 1C), samples with abnormal levels of missing data (e.g., missing rate > 30%; blue triangles), samples with high levels of non-*D. melanogaster* contamination (> 15%; green diamonds), and samples with abnormal levels of $p_N/p_S$ (>0.7% $p_N/p_S$, pink triangles). Also, a fifth category emerged from the PCA: this category describes samples that are part of the Fournier-Level et al. (2019) dataset and have a high degree of missing data. Yet, this is expected due to the low per-library coverage of the original experimental design. These samples were therefore collapsed by geographical locality (see Materials and Methods), thus improving both mean coverage (to 61X) and mean effective sample sizes ($n_e$ = 56.0).

Out of the 753 samples in the DEST 2.0 dataset, 23 were excluded due to abnormal $p_N/p_S$, and 57 due to contamination by non-*D. melanogaster* specimens, 17 due to missing data, and 136 samples were collapsed into 16 consolidated libraries. Although all DEST samples are available for users, we recommend using the resulting final high-quality dataset containing 530 samples for downstream biological analyses. In addition to these sample recommendations, DEST 2.0's companion metadata is up-to-date, and all issues reported in previous releases have been addressed (Nunez et al. 2021). Across all datasets, we discovered 4,801,077 SNPs that segregate worldwide and pass the filters of the PoolSNP calling algorithm (Kapun et al. 2020). Overall descriptions and basic subsetting of SNP statistics for DEST 2.0 are shown in Table 1. Unless stated otherwise, all of the following analyses are based on the 530 high-quality samples.
