## Supplementary material for "Footprints of worldwide adaptation in structured populations of *D. melanogaster* through the expanded DEST 2.0 genomic resource": Text S3

### Text S3. Notes on the CLL validation procedure

Prior site frequency spectrum (SFS) based demographic inference work has been unable to compare demographic models with different numbers of populations due to the dependence of model likelihoods on SFS size. We addressed this issue by introducing collapsed log-likelihood (CLL), a measure of goodness-of-fit of a demographic model that is calculated by "collapsing" dimensions of the SFSs corresponding to populations that exist in the model, but not in more parsimonious models that feature fewer populations. (see Methods of main text). Here, we describe the simulated evaluation of our CLL-based model selection strategy.

We first simulated 2D-SFSs with *msprime* (Baumdicker et al. 2022) under both the one-population and (two-population) split models for both the mainland and Americas regions with general *Drosophila melanogaster* parameters: ancestral population size of 177,344 (Kapopoulou et al. 2020), genome size of 2L, 2R, 3L, and 3R chromosomes of $1.09 \times 10^8$ (Hoskins et al. 2015), recombination rate of recombination rate $2.75 \times 10^{-3}$ cM / Mb, mutation rate of $3.265 \times 10^{-9}$ $bp^{-1}$ $generation^{-1}$ (Wang et al. 2023), and diploidy. Our recombination rate is three orders of magnitude less than what is reported in Wang et al.; this abiological choice is justified in that (1) *msprime*'s runtime is roughly linear with respect to recombination rate, making simulations of realistic recombination inhibitively slow, and (2) recombination rate has little effect on *msprime*-simulated SFSs besides controlling variance (Tran et al. 2024). Model-specific parameters (namely [$N$, $T$] for one-population and [$N_1$, $N_2$, $T$, $m$] for split) were uniformly sampled for each of $n$=40 replicates (dem_rep in the code) of each region for each model from 95% confidence intervals obtained from *moments* demographic model fits on jackknifed SFSs (see Methods). Within each replicate, we simulated 40 replicates with *msprime* (sim_rep in the code) at the same demographic parameter values, emulating the 40x jackknifing of samples within populations used on the real data. We performed demographic inference with *moments* to fit both one-population and split models to each SFS, marginalizing (i.e. summing over one axis) each 2D-SFS to yield a 1D-SFS to which the one-population can be fit and calculating collapsed log-likelihood for each fit of the split model. For each set of 40 simulation replicates at each set of demographic parameters and model, we calculated a $p$-value for the Wilcoxon signed-rank test comparing log-likelihoods of one-population model fits to collapsed log-likelihoods of split model fits for corresponding SFS simulation replicates under the null hypothesis that the one-population model does not provide a better fit to the data than the split model. Setting a significance level of $\alpha$=0.05, we were able to assess our CLL-based model selection strategy as a classifier of whether the 2D-SFS represents two distinguished populations or not. **Fig. TS3-1** shows these $p$-values, $n$=40 for each pairing of the region (from which demographic parameters were determined) and model (from which each SFS was simulated). Low $p$-values under true one-population SFSs and high $p$-values under true two-population SFSs show that our development and use of CLL for demographic model selection, at least for the mainland and Americas regions, is valid.

Future work on CLL-based model selection would examine its performance on comparing 2- and 3-population models, as we did in European populations with the Kruskal–Wallis $H$ test followed by Dunn's post hoc test instead of the Wilcoxon signed-rank test. Additionally, our strategy could be compared to simpler methods for estimating the number of populations in a sample, like the classic $F_{ST}$ threshold of 0.15 for indicating significant divergence.

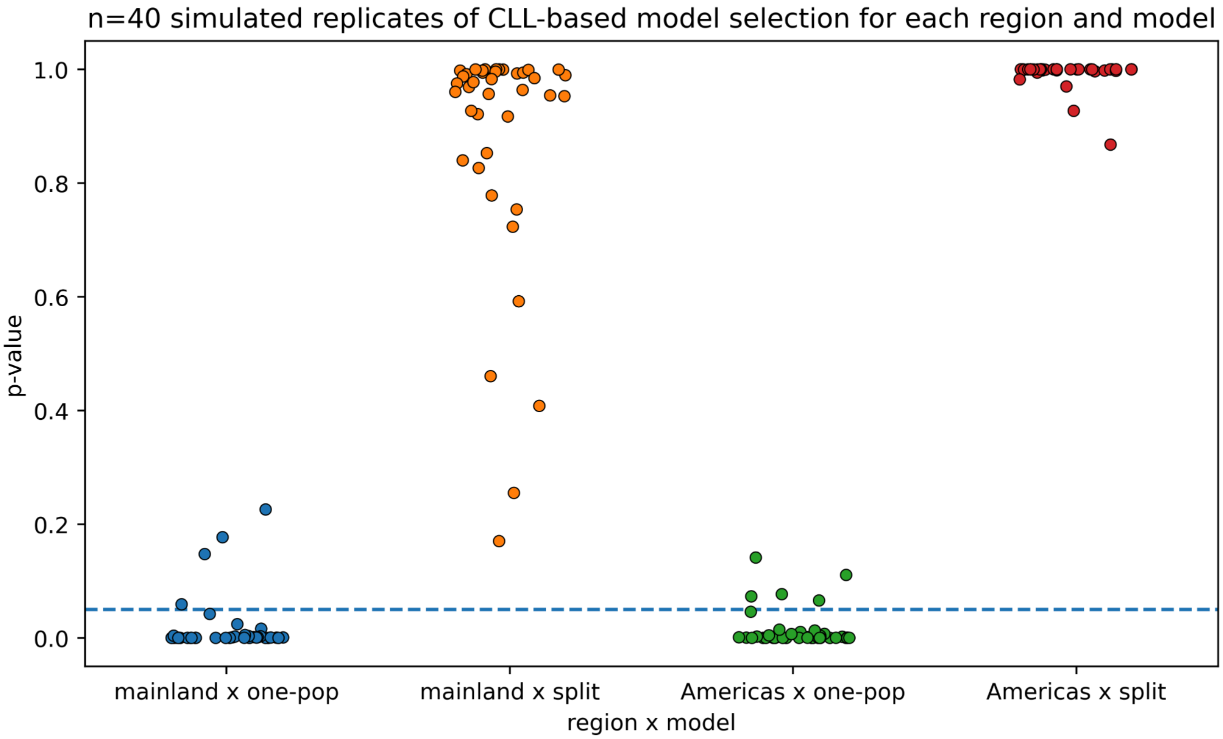


**Fig. TS3-1**: *p*-values of "$H_0$: The one-population model does not provide a better fit to the data than the (two-population) split model." from 40 simulated replicates of the model selection methodology applied to the mainland and Americas regions in the main text. Four replicates of the mainland region and five replicates of the Americas region simulated under the one-population model yield *p*-values above α=0.05, and no *p*-values for replicates simulated under the split model yield *p*-values below 0.05, so we estimate that our CLL-based model selection enjoys 100% specificity, 90% sensitivity for the mainland region, and 87.5% sensitivity for the Americas region.

---
